# A unifying photocycle model for light adaptation and temporal evolution of cation conductance in Channelrhodopsin-2

**DOI:** 10.1101/503706

**Authors:** Jens Kuhne, Johannes Vierock, Stefan Alexander Tennigkeit, Max-Aylmer Dreier, Jonas Wietek, Dennis Petersen, Konstantin Gavriljuk, Samir F. El-Mashtoly, Peter Hegemann, Klaus Gerwert

## Abstract

Although Channelrhodopsin (ChR) is a widely applied light-activated ion channel, important properties such as light-adaptation, photocurrent inactivation, and alteration of the ion selectivity during continuous illumination are not well-understood from a molecular perspective. Herein, we address these open questions using single turn-over electrophysiology, time-resolved step-scan FTIR and Raman spectroscopy of fully dark adapted ChR2. This yields a unifying parallel photocycle model explaining all data: in dark-adapted ChR2, the protonated Schiff base retinal chromophore (RSBH^+^) adopts an all-*trans*,C=N-*anti* conformation only. Upon light activation, a branching reaction into either a 13-*cis*,C=N-*anti* or a 13-*cis*,C=N-*syn* retinal conformation occurs. The *anti*-cycle features sequential H^+^ and Na^+^ conductance in a late M-like state and an N-like open-channel state. In contrast, the 13-*cis*,C=N-*syn* isomer represents a second closed-channel state identical to the long lived P_480_-state, which has been previously assigned to a late intermediate in a single photocycle model. Light excitation of P_480_ induces a parallel *syn*-photocycle with an open channel state of small conductance and high proton selectivity. E90 becomes deprotonated in P_480_ and stays deprotonated in the C=N-*syn*-cycle and we show that deprotonation of E90 and successive pore hydration are crucial for late proton conductance following light-adaptation. Parallel *anti*- and *syn*-photocycles explain inactivation and ion selectivity changes of ChR2 during continuous illumination, fostering the future rational design of optogenetic tools.

**Significance statement:** Understanding the mechanisms of photoactivated biological processes facilitates the development of new molecular tools, engineered for specific optogenetic applications, allowing the control of neuronal activity with light. Here, we use a variety of experimental and theoretical techniques to examine the precise nature of the light-activated ion channel in one of the most important molecular species used in optogenetics, channelrhodopsin-2. Existing models for the photochemical and photophysical pathway after light absorption by the molecule fail to explain many aspects of its observed behavior including the inactivation of the photocurrent under continuous illumination. We resolve this by proposing a new branched photocycle explaining electrical and photochemical channel properties and establishing the structure of intermediates during channel turnover.

## Introduction

In neuroscience, light-activated proteins are utilized to modify the membrane potential and intracellular signal transduction processes of selected cells precisely and non-invasively with light (1, 2). The first and most widely used optogenetic tool is channelrhodopsin-2 (ChR2), a light-gated ion channel from the green alga *Chlamydomonas reinhardtii* (3, 4).

ChR2 spans the membrane by seven transmembrane helices (TMH) and a retinal chromophore is covalently attached to a conserved lysine in Helix 7 *via* a protonated retinal Schiff base (RSBH^+^). Light absorption of the retinal triggers a photocycle involving spectroscopically distinguishable intermediates (Fig. 1A). After blue light excitation (λ = 470 nm) of the dark state D_470_, retinal isomerizes from all-*trans* to 13-*cis*, resulting in the red-shifted P_500_-intermediate which corresponds to K in bacteriorhodopsin (BR). Proton transfer from the RSBH^+^ to the counter ion complex leads to P_390_ (M in BR)–possibly split into an early and late P_390_ state comparable to M_1_ and M_2_ of BR (5). P_390_ is succeeded by P_520_ (N in BR) after reprotonation of the RSB. Considering the time constants of channel opening observed in electrophysiological experiments, it has been suggested that both states–the late P_390_ (M_2_) or P_520_ (N)–contribute to ion conductance of the open channel (6–8). Finally, a long-lived non-conducting state P_480_ appears after channel closing. P_480_ is considered to be the last photocycle intermediate, and from this species, D_470_ is recovered with a time constant of approximately 40 s. The unbranched photocycle is reasonably well-suited to describe a single-turnover transition starting from the dark-adapted protein, but fails to explain photocurrent changes during extended light application. During continuous illumination, photocurrents of ChR2 inactivate within milliseconds from a transient peak to stationary level and the initial peak current is recovered only after many seconds in darkness (4). Furthermore, after illumination, the current decays bi-exponentially with two distinct time constants that differ by an order of magnitude and an amplitude ratio that depends on the pre-illumination time, excitation wavelength, and membrane voltage. Thus the above mentioned single-cycle model was extended to a parallel two-cycle model comprising two closed (C_1_ and C_2_) and two open (O_1_ and O_2_) states that are populated differently in the dark and during repetitive or continuous illumination (Fig. 1B) (9, 10). Photocurrent inactivation as well as differences in conductance were explained by a higher quantum efficiency for the transition from C_1_ to O_1_ compared to that from C_2_ to O_2_, consistent with recent theoretical calculations (11).

**Figure 1:**
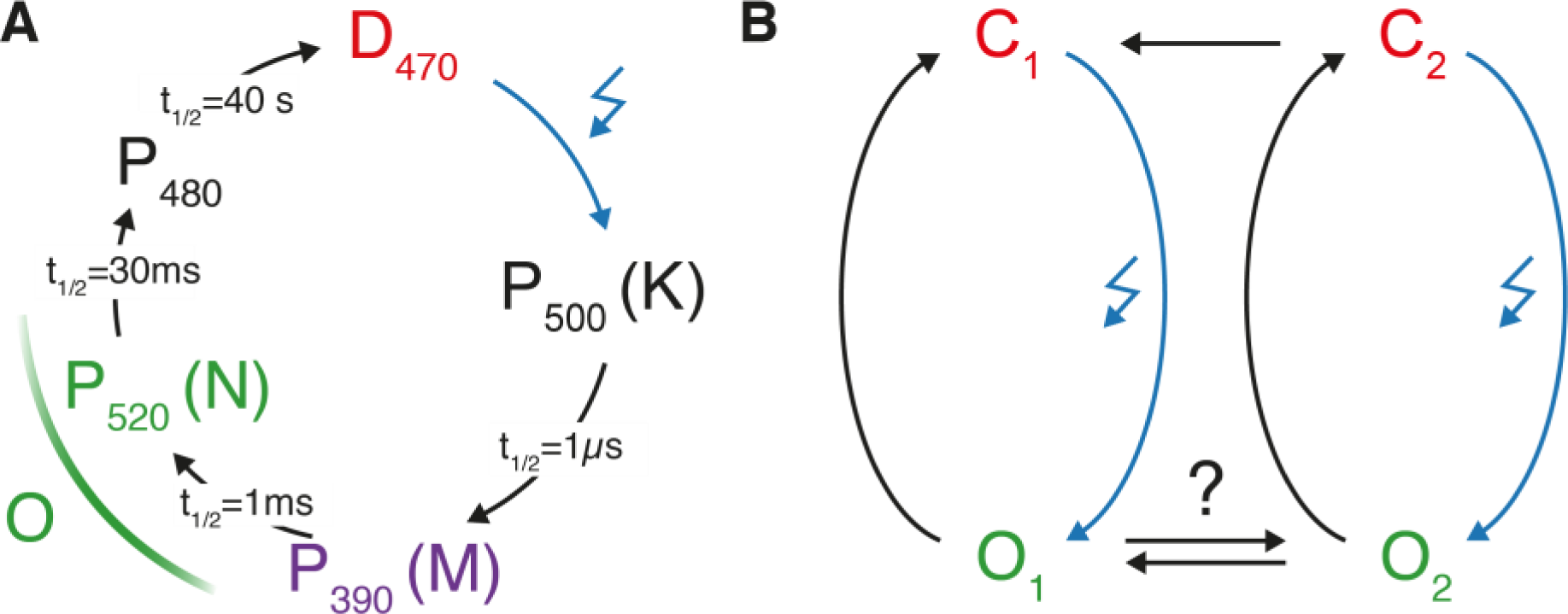
Proposed photocycle models of ChR2. (A) A non-branched photocycle model starting from the initial dark state has been most widely used to explain time resolved UV-VIS and FTIR-experiments (6, 7, 12–15). However, to explain photocurrent inactivation, a branched model (B) with two open states is required (9, 10).

Time-resolved FTIR spectroscopy has originally been established as a powerful approach for the determination of the molecular reaction mechanism of bacteriorhodopsin (16). The dark-adapted ChR2 photocycle was recorded between 50 ns and 140 s after exposure to a light pulse by step-scan and rapid-scan FTIR. These-measurements revealed an ultrafast all-*trans* to 13-*cis* isomerization and subsequent deprotonation of the RSBH^+^ in parallel with protonation of the counterion residues E123 and D253 (15). Deprotonation of D156 coincides with P_390_ depletion, which was previously considered as indicative of RSB reprotonation (14, 15). FTIR studies paired with high performance liquid chromatography (HPLC) analysis of the slow-cycling step-function variant C128T provided spectroscopic evidence for two distinct closed states with different retinal isomers (17). NMR-spectroscopic data of the ChR2 (WT) and WT-like variant H134R showed that although different closed states exist, the fully dark-adapted state (called initial dark-adapted state, IDA) of ChR2 is composed of 100% all-*trans*,C=N-*anti* retinal (18, 19). Raman experiments on ChR2-H134R revealed that illumination of the IDA at 80 K produced an apparent dark state DA_app_ containing a second retinal isomer (19). Following double isomerization around the C_13_=C_14_ and the C=N double bonds, 13-*cis*,C=N-*syn* retinal is formed, and this was proposed as the transformation step for forming the second “metastable” dark state (19). Both retinal isomers in the DA_app_ were proposed to initiate distinct photocycles, with both involving homologous P_500_-, P_390_-, P_520_-, and P_480_-like intermediates.

The central gate residue E90 is one of the key determinants of proton selectivity in ChR2 (13, 20) and related ChRs (21). During the photocycle E90, that is located in the central gate in the middle of the putative pore, is deprotonated and remains deprotonated until P_480_ decays (13–15). A late deprotonation of E90 exclusively in P_480_ was proposed in ChR2 experiments with high laser pulse repetition frequencies preventing fully dark adaptation (14). In contrast E90 deprotonation within sub-microseconds was observed in single-turnover experiments on fully dark-adapted ChR2 (15). There is a controversy between fully dark-adapted and not dark-adapted FTIR experiments on the timing of E90 deprotonation in a single photocycle model.

Here we present a unifying functional study of dark- and light-adapted ChR2 by integrating single-turnover electrical recordings and FTIR measurements on ChR2, Raman spectroscopy with ^13^C-labelled retinal and molecular dynamics simulation. The controversies observed between single-turnover experiments and recordings under continuous illumination are resolved by developing an extended model, including two parallel photocycles with 15-*anti* and 15-*syn* retinal conformations. The light-adapted 13-*cis*,15-*syn* state is the P_480_ intermediate, which was formerly assigned to the last intermediate of the *anti*-cycle in a linear photocycle model. Within the *anti-*cycle, ion conductance evolves in two subsequent steps resulting in two different conducting states of distinct ion selectivity (O_1-early_ and O_1-late_). Interestingly, E90 stays protonated in the *anti*-cycle. In contrast, the *syn*-cycle initiated by photoexcitation of P_480_, that represents the second closed state C_2_ in Fig. 1B, comprises a third slowly decaying open state O_2_ of high proton selectivity but low overall ion conductance. Conductance of O_2_ depends on deprotonation of E90 and is completely abolished in the ChR2 E90Q mutant. Our results resolve the former discrepancies. In the *anti*-photocycle E90 stays protonated and channel opening of O_1-early_ and O_1-late_, is observed, whereas in the *syn*-cycle including P_480_ E90 is deprotonated and favors proton conductance of O_2_.

## Results

### Single-turnover patch clamp recordings identify three conducting ChR2 states

In order to examine functional changes during light-adaptation of ChR2, we recorded single-turnover photocurrents in HEK293 cells following 7 ns laser excitation before and after light-adaptation. We addressed changes in ion selectivity by either reducing extracellular sodium (110 mM –> 1 mM) or proton concentration (pH_e_ 7.2 –> pH_e_ 9.0) (Fig. 2A and B).

**Figure 2:**
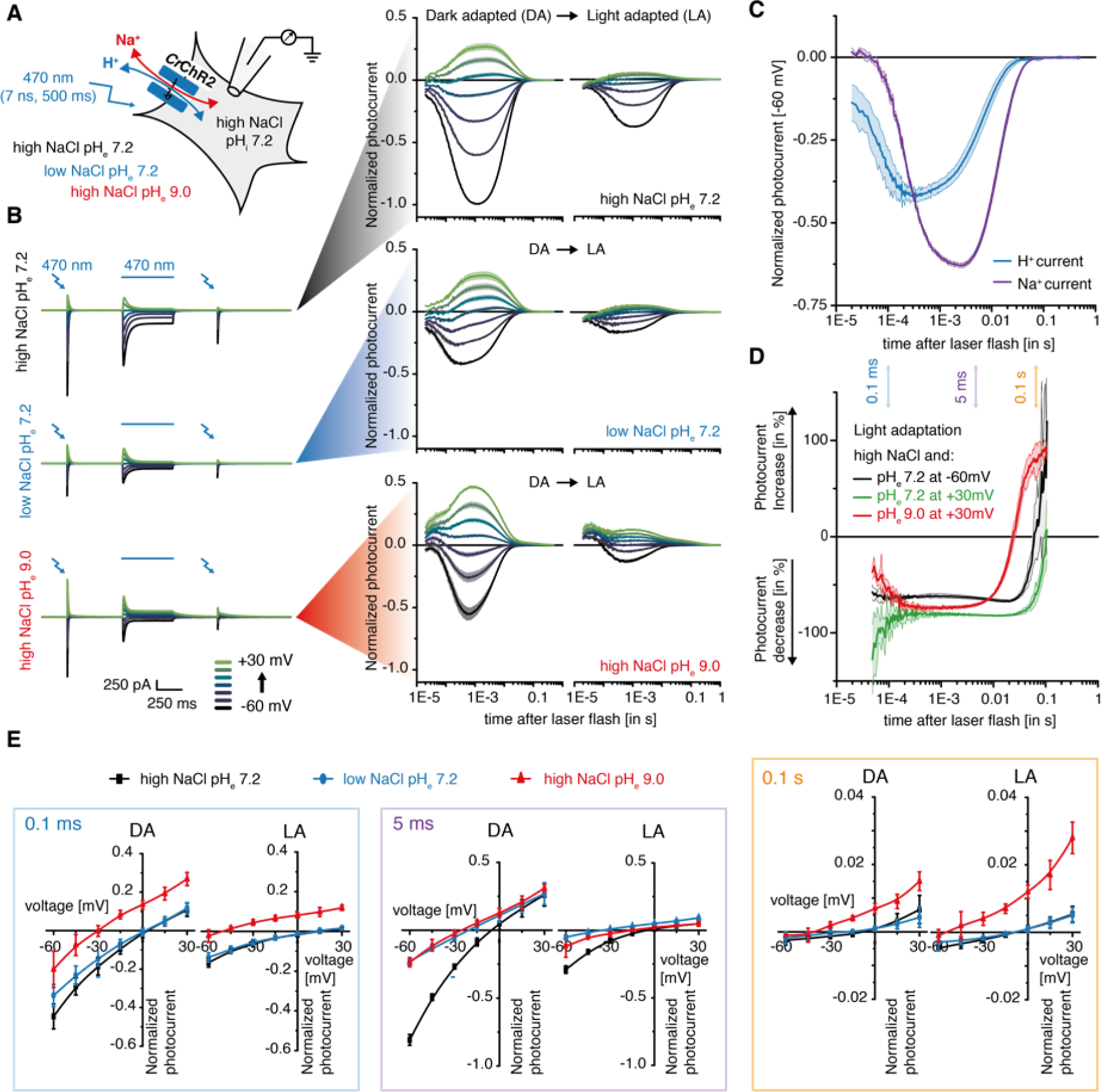
Voltage-clamp recordings in HEK293 cells of photocurrents from dark and light-adapted *ChR2* WT. (A) Experimental scheme of the whole-cell patch clamp experiment in different extracellular buffers and under different illumination conditions. (B) Left: Representative photocurrents of ChR2 with intracellular 110 mM NaCl pH_i_ 7.2 and extracellular 110 mM Na^+^ pH_e_ 7.2 (top), 1 mM Na^+^ pH_e_ 7.2 (center), 110 mM Na^+^ pH_e_ 9.0 (bottom) at different holding potentials as indicated. Photocurrents were excited before and after light-adaptation with a 470 nm 7 ns laser pulse. For light-adaptation, cells were continuously illuminated for 500 ms with 470 nm light. Right: Normalized, log-binned and averaged photocurrents of the dark-adapted (DA) or light-adapted (LA) protein (mean ± SEM, n = 5–8). (C) Time evolution of estimated proton and sodium fluxes in the dark-adapted protein at −60 mV either directly measured in extracellular 1 mM Na^+^ pH 7.2 (H^+^ current) or calculated by subtraction of proton fluxes from combined inward flux of sodium and protons measured in symmetric conditions (Na^+^ current, I(110 mM Na^+^ pH 7.2)-I(1 mM Na^+^ pH 7.2), mean ± SEM, n = 7). (D) Relative photocurrent changes upon light-adaptation at different extracellular voltages and pH_e_ (I(LA)-I(DA))/I(DA), mean ± SEM, n = 5–8). (E) Current-voltage dependency of normalized photocurrents at 0.1 ms (left), 5 ms (center) and 100 ms (right) after excitation in different extracellular buffer compositions before (DA) and after (LA) light-adaptation (mean ± SD, n = 5–8).

Under symmetric sodium and proton concentrations, the dark-adapted ChR2 pore opens bi-exponentially with two almost voltage-independent time constants (150 µs and 2.5 ms). The photocurrents decline with a dominant voltage-dependent time constant of 10–22 ms and a second, minor, slow time constant of 70–220 ms (Fig. 2A and B, top), in general agreement with previous reports (8). Decreasing extracellular Na^+^ not only reduces inward directed photocurrent amplitudes, but also affects the temporal evolution of inward currents (Fig. 2A, center). Whereas inward currents in low extracellular Na^+^ are predominantly carried by protons (H^+^ flux), inward currents under symmetric conditions are mediated by both H^+^ and Na^+^ ions. Subtraction of photocurrents at high and low Na^+^ at pH 7.2 allows an approximation of the pure inward Na^+^ flux (Fig. 2C). Strikingly, the proton flux peaks as early as 300 µs after excitation, significantly earlier than Na^+^ flux (2.5 ms). This observation is indicative of two open states with distinct ion selectivity following single excitation of dark-adapted ChR2.

During continuous illumination, photocurrents peak within milliseconds (dependent on the light intensity) and subsequently decline to a stationary level. Inactivation is more pronounced at positive voltages, contributing to the increased inward rectification of stationary photocurrents compared to the initial peak current (4). After light-adaptation, laser pulse-induced photocurrents are significantly reduced in amplitude. But still the photocurrent rises and decays bi-exponentially, reaching a maximal amplitude at the same post-flash time point as photocurrents in the dark-adapted protein (Fig. 2B). The relative photocurrent change at different time points after excitation shows homogeneous photocurrent reduction of 60–80% between 0.2 and 10 ms (Fig. 2D). However, notably, relative photocurrent changes differ at early time points (< 200 µs) and during the slow photocurrent decline (after 50–100 ms), indicating different open state conformations in the early and late stages of the photocycle of light- and dark-adapted ChR2. In particular, at 100 ms after the laser excitation, the photocurrents even increase in amplitude for light-adapted ChR2 (especially at pH 9.0), indicative of an additional slowly-decaying open state. In summary, there are at least three different conductive states, O_1-early_ and O_1-late_ in the dark-adapted photocycle and O_2_ after light-adaptation.

The proton versus sodium selectivity of all three open states is analyzed at three different time points after the actinic laser pulses (0.1 ms, 5 ms and 100 ms; Fig. 2E). Although the photocurrents at 0.1 ms and 100 ms are barely distinguishable in high and low extracellular Na^+^ in either the dark- or the light-adapted protein, reduction of extracellular proton concentration causes a strong shift in reversal potential and an increase in outward-directed photocurrent amplitudes that is even more pronounced in the light-adapted than in the dark-adapted channel. In contrast, for photocurrents 5 ms after excitation, both ionic changes–reduction in extracellular sodium or proton concentration–shift the reversal potential and decrease the inward photocurrent amplitude. We conclude that after channel opening, the short lived highly proton-selective O_1-early_ is followed by the more Na^+^-selective, but still highly proton-permeable O_1-late_. After multiphoton excitation and light-adaptation, the contribution of O_1-early_ and O_1-late_ decreases in favor of the third highly proton-selective O_2_, which, although small in amplitude, significantly contributes to stationary photocurrents at alkaline pH due to its long lifetime.

### Single-turnover time-resolved FTIR measurements reveal a splitting of the photocycle after light-activation of fully dark-adapted ChR2

The single-turnover electrophysiology data recall that dark adaptation and light-adaptation need to be compared thoroughly for the correct interpretation of time resolved measurements. However, most time resolved spectroscopy studies are performed with barely dark-adapted samples at rather high repetition rates in order to avoid long measurement times. To elucidate the underlying molecular mechanism of the observed channel gating transitions and different ion conductance, we performed single-turnover time-resolved FTIR measurements of the fully dark-adapted ChR2 WT-like H134R variant with a time resolution of 50 ns over nine orders of magnitude (Fig. 3A) similar to our data from 2015 (15). The ChR2 WT-like H134R mutant shows higher protein expression in *Pichia pastoris* compared to the wild-type protein and has been used for the examination of light-adaptation before (19). Electrical properties and photocycle kinetics are comparable although slightly slower than the ones of the WT protein (22), and the same IR-bands are observed in WT and in H134R. However, some crucial infrared marker bands are more pronounced in H134R which simplifies the presentation of the dataset. Dark adaptation of D_470_ was achieved by long dark periods of 140 s between pulsed excitation (T = 15 °C), that increased the step-scan measurement time to about 4 weeks, whereas light-adapted samples take few hours only. The unresolved early appearance of the marker band at 1188 cm^−1^, indicates the all-*trans* to 13-*cis*,C=N-*anti* isomerization, because it represents the C_14_-C_15_ stretching vibration of 13-*cis* retinal as originally assigned in bacteriorhodopsin by site-specific isotopic labeling (23). The decay of the 1188 cm^−1^ marker band within a microsecond (green time trace in Fig. 3A) indicates the formation of the M-like P_390_ intermediate with a deprotonated RSB (15). As in other microbial rhodopsins, the subsequent rise and decay of the N-like P_520_ intermediate with a reprotonated Schiff-base can be monitored by its reappearance and the decay reflects formation of the all-*trans* isomer on a few-millisecond timescale. Comparing the time course of this marker band (1188 cm^−1^) with the single-turnover electrical measurements, we can assign the described conducting states O_1-early_ and O_1-late_ to the late part of P_390_ (M_2_) and P_520_ (N) respectively, which is in line with earlier reports on the WT protein (8). Due to these similarities and the abundance of spectroscopic data on BR, we decided to name the ChR2 intermediates as follows: P_500_^K^, P_390a_^M1^, P_390b_^M2^, and P_520_^N^. Global fitting of the whole dataset (solid lines) describes the data adequately. The apparent rate constants of H134R are similar with earlier reports for the dark-adapted ChR2 WT (15).

**Figure 3:**
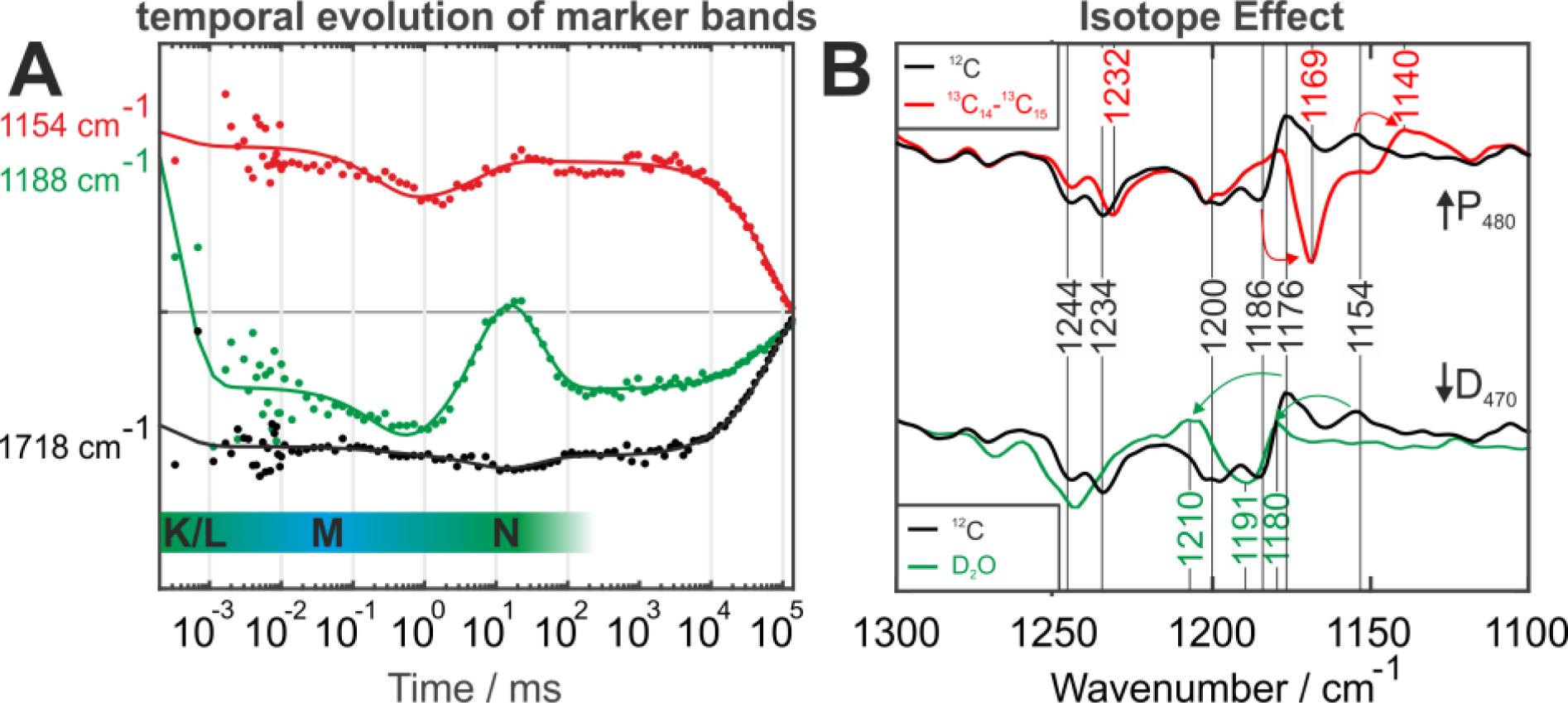
FTIR measurements on H134R and WT. (A) Kinetic transients of the marker bands in WT-like H134R variant recorded by step-scan FTIR. The P_480_ C=N-*syn* (red) and E90 (black) marker bands, 1154 cm^−1^ and 1718 cm^−1^ respectively, are observed not time-resolved at the very beginning of the reaction. Their decay occurs with the decay of P_480_ (t_1/2_ = 40 s). In parallel, the D_470_ → P_500_^K^ → P_390_^M^ → P_520_^N^ reaction is monitored by the marker band for protonated 13-*cis*,C=N-*anti* retinal at 1188 cm^−1^ (green). The continuous lines are the result of a global fit analysis using five rate constants that sufficiently describe the dataset. (B) Comparison of unlabeled (black), ^13^C_14_-^13^C_15_ labeled (red) and unlabeled but deuterated (green) WT-samples in the P_480_-D_470_ difference spectrum. The marker band at 1188 cm^−1^ from A is not seen at this late photocycle intermediate. The red arrows indicate the isotope induced downshifts of the C_14_-C_15_ stretching vibration at 1186 cm^−1^ in D_470_ and 1154 cm^−1^ in P_480_. The green arrows denote the large upshift of the C_14_-C_15_-stretching vibration at 1154 cm^−1^ induced by deuteration to 1180 cm^−1^ indicating the *syn*-conformation. Also, the C_10_-C_11_ stretching vibration at 1176 cm^−1^ is upshifted. The large upshift of the P_480_ bands indicates a C=N-*syn* conformation of the retinal in P_480_. In contrast the C_14_-C_15_-stretching vibration at 1186 cm^−1^ in D_470_ is only slightly upshifted in D_2_O to 1191 cm^−1^ indicating a trans conformation. For more details see Figs. S1-S8, Notes 1 and 3–5 in the SI Text, and Table S1.

### Light-induced splitting in 13-*cis*,C=N-*anti* and 13-*cis*,C=N-*syn* RSBH^+^ conformations

Interestingly, an additional retinal band at an unusual low wavenumber, 1154 cm^−1^, appears faster than the time resolution of the instrument parallel to the 1188 cm^−1^ 13-*cis*,C=N-*anti*-marker band. Because the low wavenumber band is more pronounced in H134R than in WT (Fig. S1), we discuss the data here for the reader on the mutant although the results are also valid for the WT. The 1154 cm^−1^ band persists from nanoseconds to seconds after a single pulse of excitation light (Fig. 3A). The band is assigned here to the C_14_-C_15_-stretching vibration of retinal because of the characteristic 14 cm^−1^ downshift upon retinal ^13^C_14_-^13^C_15_ carbon-specific labeling (Fig. 3B upper part). The band assignment is confirmed by additional Raman experiments (Fig. S6). The 22 cm^−1^ upshift of the C_14_-C_15_ band in D_2_O indicates a 13-*cis*,C=N-*syn* conformation (Fig. 3B lower part). In 13-*cis*,C=N-*syn* retinal, the C_14_-C_15_-stretching vibration is strongly coupled to the N-H-bending vibration, which is decoupled in D_2_O (N-D) and results in a deuteration induced large upshift in the *syn* but not in the *anti*-conformation (24, 25). Therefore, the band at 1154 cm^−1^ represents a 13-*cis*,C=N-*syn* marker band. The band assignments are nicely confirmed by Raman experiments shown in the supplement in more detail for P_480_ (Figs. S1-S6, Notes 1 and 3–5 in the SI Text, and Table S1).

The negative difference bands in Fig. 3B reflect vibrations of dark-adapted ChR2 WT (D_470_). The negative band at 1186 cm^−1^ is also assigned to the C_14_-C_15_-stretching vibration of D_470_ because of the characteristic isotope downshift. From the additional analysis of the D_470_ Raman spectrum (Supplemental Note 3 and 4) we conclude that the retinal of dark-adapted ChR2 is in an 100% all-*trans*,C=N-*anti* conformation, in agreement with NMR data (18, 19). For a detailed band assignment of the P_480_ and D_470_ vibrational spectra and retinal conformations, the reader is referred to Figs. S1-S7, Notes 1 and 3–5 in the SI Text, and Table S1.

The parallel but temporally unresolved appearance of the bands at 1188 cm^−1^ and 1154 cm^−1^ in single-turnover experiments in Fig. 3A indicates that light-absorption induces parallel isomerization of all-*trans,C=N-anti* retinal in D_470_ into either a 13-*cis*,C=N-*anti* or a 13-cis,C=N-*syn* conformation. The splitting ratio into parallel *syn-* and *anti-*pathways can be estimated as 1:1 under our measurement conditions (Fig. S3).

Considering that the 13-*cis,*C=N-*syn* isomerization occurs in parallel to the 13-*cis,*C=N-*anti* isomerization, we conclude that the 13-*cis*,C=N-*syn* retinal conformation observed in P_480_ is therefore not the last intermediate of the 13-*cis*,C=N-*anti* photocycle, as proposed in the single-cycle model. P_480_ reflects a long-lived 13-*cis*,C=N-*syn* state which appears in parallel to 13-*cis*,C=N-*anti* instantaneously.

The conclusion that the P_480_ is not the last intermediate of the 13-*cis*, C=N-*anti* single photocycle but appears in parallel as light-adapted state is furthermore strongly supported by detailed Raman experiments as described in in the supplement. The Raman results are in agreement with former Raman studies on light-adaption (19). Upon complete light-adaptation of ChR2 due to long illumination periods an apparent dark state D_app_ evolves. It is composed of a 40:60 mixture of the all-*trans*,C=N-*anti* species D_470_ and the 13-*cis,*C=N-*syn* species in P_480_ (Supplementary Fig. S3). The bands observed in the Raman spectra of D_470_ and P_480_ correlate with retinal bands seen in the infrared difference spectra in Fig. 3B and agree with the published Raman spectra of the all-*trans*,C=N-*anti* and 13-*cis*,C=N-*syn*-bands of the D_app_ state (19) (Supplementary Note 3). The Raman data confirm the ultrafast C=N-*syn* formation in P_480_ as seen in Fig. 3A at 1154 cm^−1^.

### E90 deprotonates upon 13-*cis,*C=N-*syn* formation

The E90-deprotonation marker band (1718 cm^−1^) (15) and the C=N-*syn* marker band (1154 cm^−1^; Fig. 3A) appear instantaneously, not time resolved. Both marker bands persist alongside the dark-adapted *anti-*photocycle intermediates (P_500_^K^, P_390_^M^ and P_520_^N^). Both decay with a slow half-life of approximately 40 s. We therefore conclude that E90 remains deprotonated during the entire C=N-*syn* pathway. However, E90 does not deprotonate in the *anti*-cycle. This resolves the former discrepancies between Lórenz-Fonfría et al. (14) and Kuhne et al. (15). In agreement with the former findings E90 deprotonates in P_480_, but this intermediate and E90 deprotonation appears not late in the last intermediate in a linear photocycle (Lórenz-Fonfría (14)), but much faster with appearance of P_480_ during light-adaptation (Kuhne (15)). In addition, E90 deprotonation appears to be closely connected with helix hydration, as indicated by the helix hydration marker bands (1662 cm^−1^ in D_470_ and 1650 cm^−1^ in P_480_) that were assigned in an earlier study, but for only for P_480_ (8). Interestingly, the same P_480_ hydration marker bands that are present in the WT were no longer observed in the mutants E90Q, E123T, and K93S that prevent E90 deprotonation (Fig. S1 and Note 2 in the SI Text). Therefore E90 deprotonation seems to be induced by the all-*trans*, C=N-*anti* → 13-*cis,*C=N-*syn* isomerization and modulates the water influx in P_480_.

### Excitation of P_480_ induces a parallel photocycle

After light-adaptation both D_470_ and P_480_ serve as parent states for parallel photocycles. At flash frequencies of 0.2 Hz that do not allow sufficient dark state recovery, a bi-exponential decay with a fast (*t*_1/2_ = 30 ms) and slow process (*t*_1/2_ = 250 ms) was observed. The corresponding amplitude spectra are shown in Fig. 4C (see also Fig. S8 and Note 6 in the SI Text). The 30 ms amplitude spectrum exhibits positive UV/VIS bands at 380 nm and 520 nm (Fig. 4B), indicative of a mixture of P_390_^M^ and P_520_^N^ intermediates in the C=N-*anti*-cycle. In contrast, if the sample is sufficiently dark-adapted (0.005 Hz flash repetition rate) the 250-ms component is no longer observed. We therefore conclude that the slow time constant reflects the conducting state of the *cis*-cycle (Fig. 4A). Because the *t*_1/2_ = 30 ms process reflects an apparent but not an intrinsic rate constant, the decay of both intermediates, which follows different intrinsic rate constants, is described by the integrated apparent rate.

**Figure 4:**
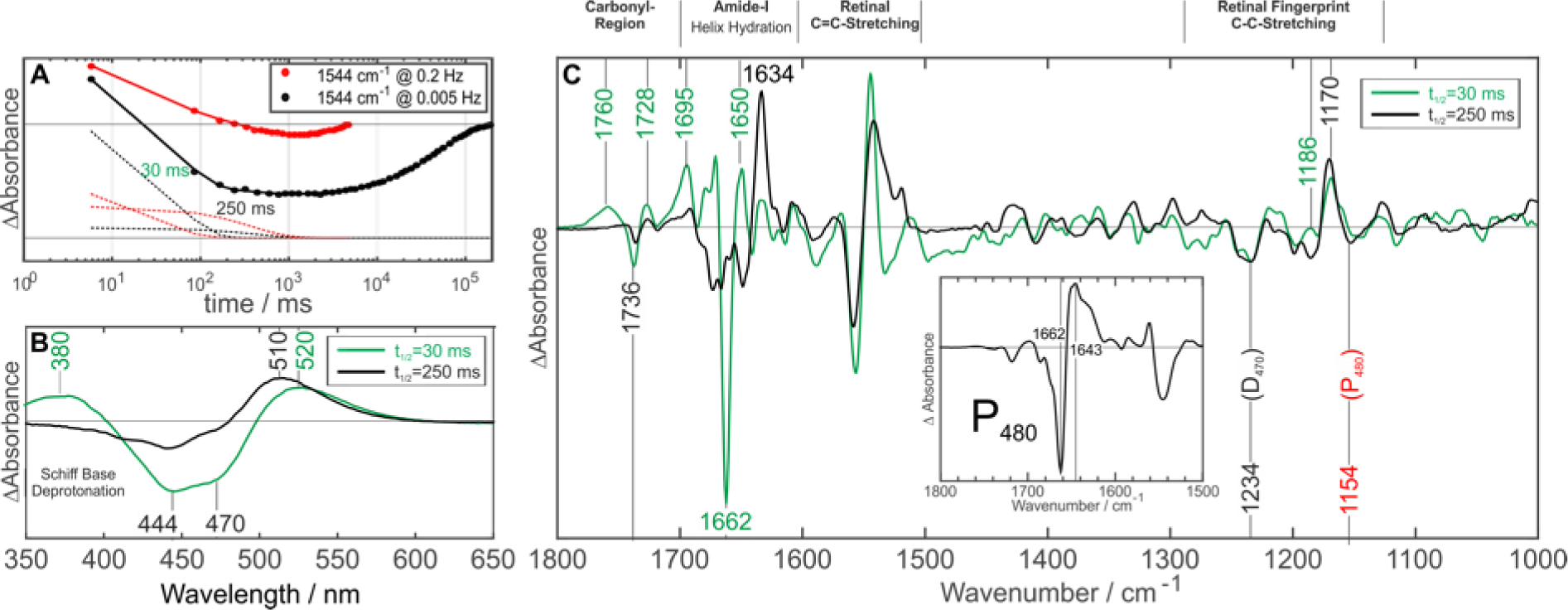
Kinetic behavior of ChR2-WT at different laser pulse repetition rates. (A) Time-evolution of the Amide-I band at 1544 cm^−1^ at low (0.005 Hz, black) and high (0.2 Hz, red) pulse repetition frequency. Upon higher pulse repetition frequency, the decay switches from mono-to bi-exponential. (B) Comparison of O_1_ (green) and O_2_ (black) decay associated UV/VIS amplitude spectra. The two positive bands at 380 nm and 520 nm in O_1_ (green) indicate a mixture of P_390_^M2^ (O_1-early_) and P_520_^N^ (O_1-late_). No evidence for RSBH^+^ deprotonation is visible in the slow component (O_2_). (C) The same processes as monitored by FTIR.

The amplitude spectrum of the 250 ms apparent rate constant indicates decaying intermediates of the *syn*-cycle. As there is only a positive band at 520 nm, it was designated the P*_520_^N^ intermediate to distinguish it from the P_520_^N^ of the *anti*-cycle. The P*_520_^N^ kinetics fit well to the slow photocurrent component O_2_ and is assigned to it in the following. In contrast, no P_390_^M^-like intermediate is observed in the C=N-*syn*-cycle (Fig. 4B).

### FTIR amplitude spectra indicate structural differences of O_1_ **and O**_2_

The O_1_- and O_2_- FTIR-amplitude spectra of the 30 ms and 250 ms apparent rates are shown in Fig. 4C. Both decay-associated amplitude spectra exhibit negative D_470_ marker bands (Fig. 4C, Figs. S8 and S10) indicating a direct transition from the *anti-* and the *syn-*photocycle into the all-*trans*,C=N-*anti* configuration of D_470_.

The *t*_1/2_ = 30 ms FTIR decay associated amplitude spectrum of the C=N-*anti*-cycle exhibits carbonyl bands at 1760 cm^−1^ (+)/1736 cm^−1^ (-) and 1728 cm^−1^ and 1695 cm^−1^ which were assigned to protonation of the counter ion D253 (15, 26) (1728 cm^−1^) and deprotonation of D156 (1736 cm^−1^). Furthermore, the helix hydration marker bands at 1662 cm^−1^ (-)/1650 cm^−1^ are present, which are now assigned to both O_1-early_ and O_1-late_. In the *t*_1/2_ = 250 ms decay-associated amplitude spectrum, all carbonyl bands are strongly reduced, including the bands of the SB proton acceptor D253 (at 1728 cm^−1^) as well as D156 (1736 cm^−1^), which has been proposed to be the RSB reprotonation donor (14). Because the Schiff base deprotonation is not observed the corresponding counterion D253 protonation is not seen as well. Also, reprotonation of D156 (1736 cm^−1^ band) is no longer observed. The negative P_480_-band (1154 cm^−1^) in the *t*_1/2_ = 250 ms decay-associated amplitude spectrum indicates an additional O_2_ → P_480_ back reaction within the *syn*-photocycle (Fig. S8 and S10 and Note 6 in the SI Text).

### Isomerization of retinal leads to a rearrangement of the central gate

To visualize the structural changes within the protein, we performed molecular dynamics simulations based on the recently published crystal structure of ChR2 WT (PDB-ID: 6eid) (27). The structure of ChR2 is highly similar to the structure of the ChR1-like chimera C1C2, but the in contrast to the latter, in the ChR2 structural model E90 is already downward oriented in the dark adapted state and at the same position as observed for the C1C2 chimera structural model after isomerization (15, 28). Within our here presented simulations of the ChR2 WT structure the retinal isomerization was changed from the dark-adapted all-*trans*,C=N-*anti* conformation (see Fig. 5C left panel) to either a 13-*cis,*C=N-*anti* single isomerization (see Supplementary Note 7, Fig. S13C+D) or a 13-*cis,*C=N-*syn* double isomerization (see Fig. 5C right panel, Fig. S13B). The observed changes in hydrogen bond interaction and water distribution are shown in Fig. 5B+C and Figs. S12-S14. Noteworthy, the single isomerization induces an upward orientation of the RSB proton, whereas, in the double isomerization, the position of the RSB proton is only slightly changed (13, 19) (Fig. 5A). Starting from the WT structure E90 keeps its initial downward orientation in the dark-adapted state (Fig. 5B+C). Very recently a more advanced method to perform such isomerization simulations was introduced by Ardevol and Hummer (28). They simulated a homology model of ChR2 based on the C1C2 chimera crystal structure (PDB-ID: 3ug9 (29)) and obtained a downward flip of the initially upward orientated E90. We have already observed the same downward movement in our model based on the same crystal structure using a classical approach (15). This proves that even our classical approach predicts correctly alterations of hydrogen bond pattern of E90 due to retinal isomerization. It seems that E90 is trapped in a local minimum in both models but finds the correct position for ChR2 (as observed in the PDB-ID: 6eid crystal structure (27)) after disturbance by isomerization.

**Figure 5:**
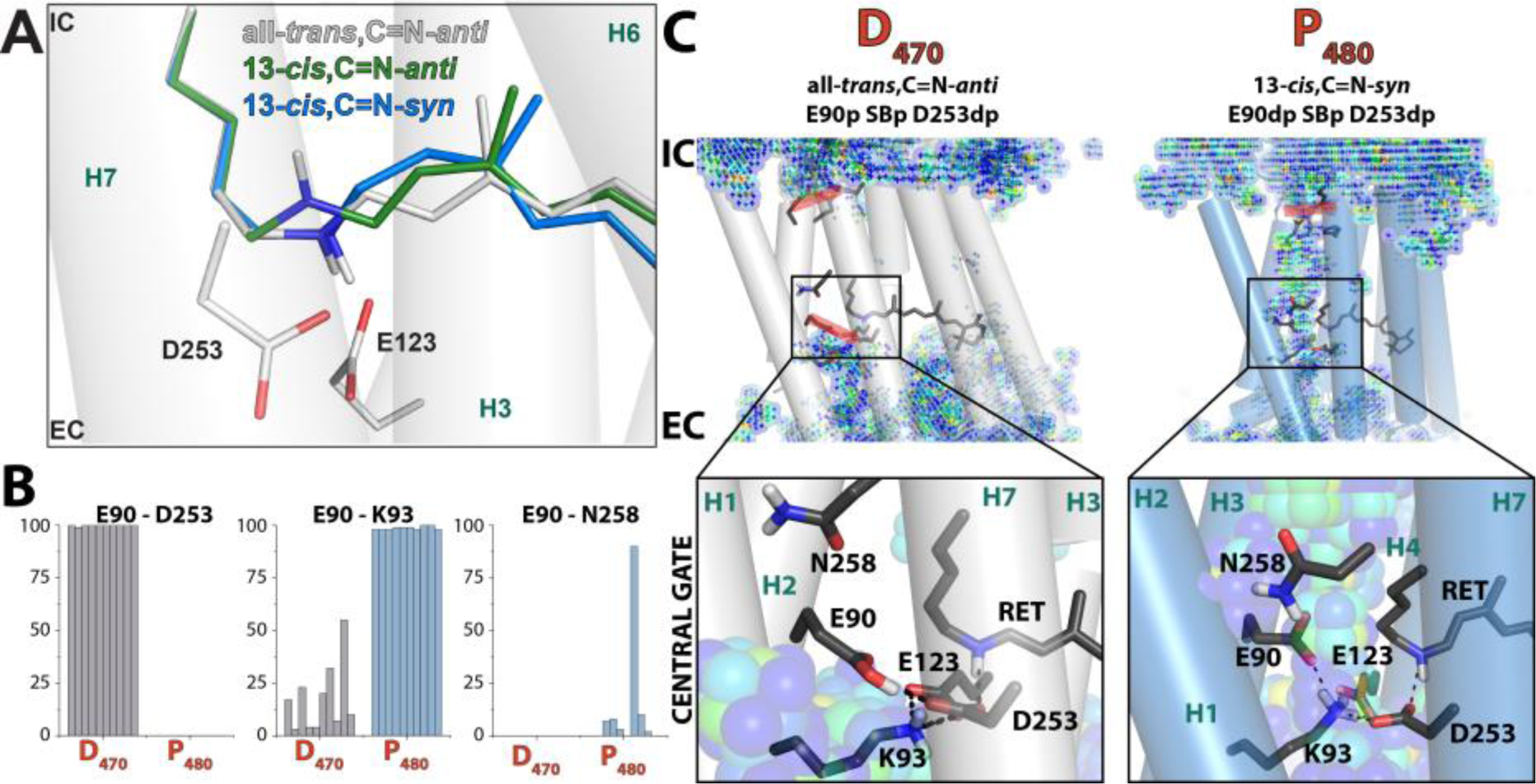
Retinal conformations and formation of P_480_. (A) A modeled representation of the calculated retinal configurations. The Schiff base orientation in the D_470_ structure all-*trans*,C=N-*anti* (gray), and modeled 13-*cis*,C=N-*anti* (green) and 13-*cis*,C=N-*syn* (blue) retinal structures are shown. (B) Overview of E90 hydrogen bond pattern for 5 independent simulations with two monomers forming one dimer based on the ChR2 WT crystal structure (PDB-ID 6eid (27)). Bars indicate the frequency of the respective hydrogen bond in percent during the 100 ns simulation. (C) The representative structure of the simulations is depicted. The left panel shows the D_470_ dark state. The right panel shows the structures after all-*trans*,C=N-*anti* → 13-*cis*,C=N-*syn* double isomerization and E90 deprotonation (P_480_).

Following 13-*cis,*C=N-*syn* double isomerization Helix 2 and 7 stay connected via E90 and D253 as long as E90 remains protonated (Fig. S12). Deprotonation of E90 leads to an alternative contact between E90 and K93 (Figs. 5B+C, S12), that opens the central gate and results in an influx of water molecules into the pore (Fig. 5C). This water influx agrees to the former channel opening due to E90 deprotonation observed in the EHT model (15) but for the 13-*cis*,C=N-*anti* conformation. We attribute E90 deprotonation and pore hydration now to the light-adapted closed state P_480_. In this light-adapted state the inner gate still remains closed and ion permeation is hindered in agreement with the electrophysiology results (Fig. S14). As 13-*cis*,C=N-*syn* isomerization accumulates during light-adaptation, we could attribute E90 deprotonation and pore hydration to the lightadapted closed state P_480_.

### Deprotonation of E90 is essential for proton conductance of O_2_ following light-adaptation

As we have shown that the deprotonation of E90 is responsible for pore hydration in the light-adapted dark state P_480_, an important role of E90 deprotonation for channel conductance in the *syn*-cycle appeared likely from our model. Consequently we mutated E90 to glutamine and analyzed photocurrent changes before and after light-adaptation (Fig. 6A, Fig. S11). In general agreement with previous results of steady state measurements (13, 30), the E90Q mutation reduces proton conductance of O_1-late_ of the *trans*-cycle. Accordingly, upon reduction of extracellular sodium photocurrent amplitudes are more decreased compared to the wild-type (Fig. 6B) and the reversal potential shifts positive 2 ms after excitation (Fig. 6C, Fig. S11C). In addition to the effect on the *anti*-cycle, the E90Q mutation also completely abolishes the late photocurrent increase upon light-adaptation, which was observed in the WT channel (Fig. S11D). Instead, slow photocurrents are reduced in the E90Q mutant following continuous illumination (Fig. 6D and E). This indicates, that E90 facilitates proton conductance of O_2_ by deprotonation, rendering it completely impermeable in the E90Q mutant. The results on the E90Q mutation validate our photocycle model with a parallel *syn*-photocycle that involves E90 deprotonation and populates during light-adaption.

**Figure 6:**
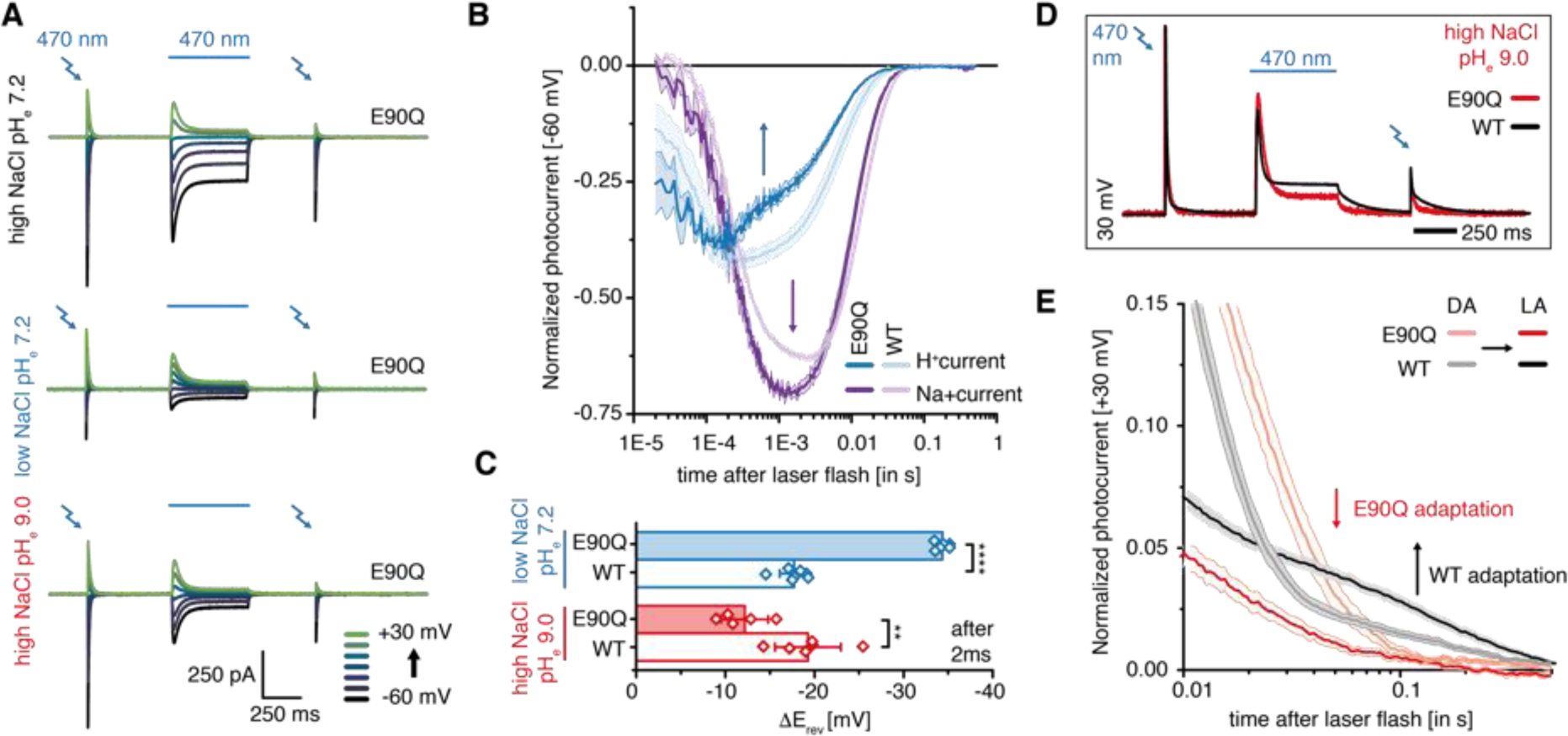
Proton and sodium conductance of the dark- and light-adapted *ChR2* mutant E90Q. (A) Representative photocurrents of ChR2-E90Q with intracellular 110 mM NaCl pH_i_ 7.2 and extracellular 110 mM Na^+^ pH_e_ 7.2 (top), 1 mM Na^+^ pH_e_ 7.2 (center), 110 mM Na^+^ pH_e_ 9.0 (bottom) at different holding voltages as indicated. Photocurrents were excited before and after light-adaptation by 7 ns laser pulses of 470 nm wavelength light. For light-adaptation, cells were illuminated for 500 ms with continuous 470 nm light. (B) Time evolution of estimated proton and sodium fluxes in the dark-adapted protein at −60 mV either directly measured in extracellular 1 mM Na^+^ pH 7.2 (“H^+^ current”) or calculated by subtraction of proton fluxes from combined inward flux of sodium and protons measured in symmetric conditions (“Na^+^ current”, I(110 mM Na^+^ pH 7.2)-I(1 mM Na^+^ pH 7.2), Mean ± SE, WT n = 7, E90Q n = 6). (C) Reversal potential shift 2 ms after laser light excitation of the dark-adapted protein upon reduction of extracellular sodium (110 mM NaCl –> 1 mM NaCl) or proton (pH_e_ 7.2 –> pH_e_ 9.0) concentration (Mean ± SD, E90Q n = 5–6, WT n = 6–7, corrected for liquid junction potentials). (D) Equally scaled representative photocurrents of *ChR2* WT and E90Q at +30 mV and extracellular 110 mM Na^+^, pH_e_ 9.0. (E) Normalized, log-binned and averaged photocurrents of the dark-adapted (DA) or light-adapted (LA) WT and E90Q at +30 mV and extracellular 110 mM Na^+^, pH_e_ 9.0 (mean ± SEM, WT n = 6, E90Q n = 5).

## Discussion

In the present work, we combined single laser pulses and continuous or repetitive illumination and analyzed the fully dark- and light-adapted ChR2 in single-turnover electrophysiological recordings, time-resolved FTIR and resonance Raman spectroscopic measurements, complemented by MD-simulations. We verified early branching into two parallel photocycles with distinct retinal isomerization and alternative configurations of the central gate and elaborated a unifying photocycle model shown in Fig. 7 that addresses light-adaptation and temporal changes in cation conductance on a functional and molecular level.

**Figure 7:**
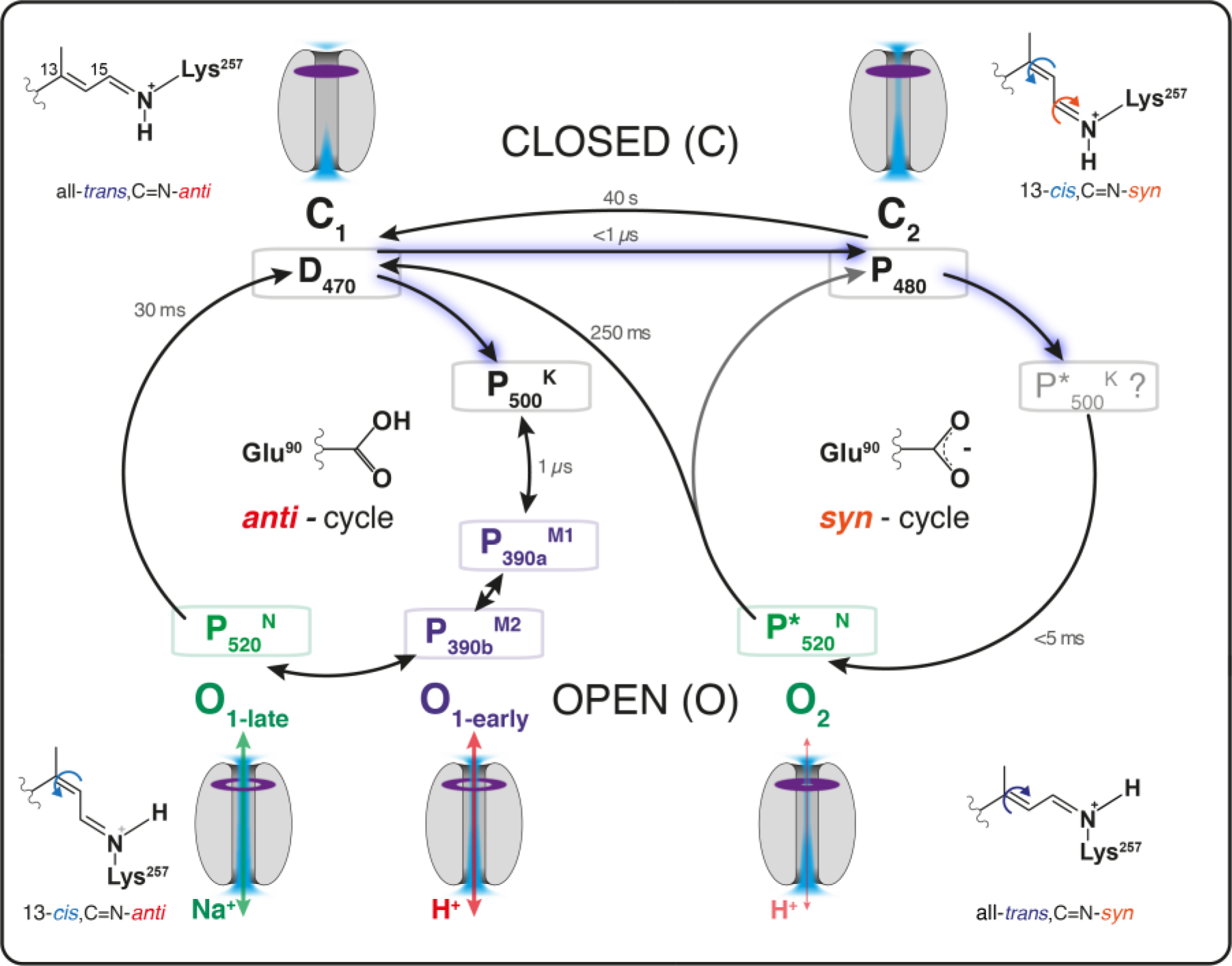
Unifying photocycle model: D_470_ and P_480_ represent the closed states C_1_ and C_2_ of Fig. 1B. D_470_ is the state with all-*trans*,C=N-*anti* retinal which is populated to almost 100% in fully dark-adapted ChR2. Upon light activation, two pathways are observed. Path 1: all-*trans*,C=N-*anti* → 13-*cis,*C=N-*anti* isomerization initiates the *anti*-photocycle, with the K-like P_500_^K^ that converts into M-like P_390_^M^ and N-like P_520_^N^ assigned to the open states (O_1-early_) and (O_1-late_). Path 2: all-*trans*,C=N-*anti* → 13-*cis*,C=N-*syn* isomerization leads to P_480_-formation with deprotonated E90. The non-conducting state P_480_ slowly relaxes back to D_470_. At high flash frequencies or under continuous illumination, P_480_ (C_2_) accumulates. The light-adapted apparent dark state represents a mixture of D_470_ and P_480_ (C_1_ + C_2_). The photoproduct of the photo-reactive P_480_ (C_2_) is the N–like P*_520_^N^, which is regarded as the open state O_2_. Due to its relatively long decay time P*_520_^N^ accumulates under continuous illumination with bright light as commonly used in electrophysiology. During such conditions both P_480_ and P*_520_^N^ contribute to the inactivation of ChR2 as they accumulate at the expense of the highly conductive O_1_ and its parent state D_470_. E90 remains protonated during the *anti*-cycle and deprotonated during the *syn*-cycle.

Following longer dark periods the “initial dark state” IDA comprises only D_470_ containing 100% all-*trans*,C=N-*anti* retinal, which is in agreement with previous reports (18, 19). The inner gate and the central gates are closed and interhelical hydrogen bonding of D253 with the RSBH and of the protonated E90 with D253 (28) prevent the invasion of water molecules from the extracellular bulk phase. After illumination of D_470_ (C_1_), early branching of the photocycle due to an alternative retinal double isomerization occurs.

In the reaction path starting from D_470_ (designated the *anti*-cycle) all-*trans*,C=N-*anti* retinal isomerizes to 13-*cis,*C=N-*anti* leading to P_500_^K^, deprotonation of the RSBH^+^ in P_390_^M^ and reprotonation of the RSB in P_520_^N^, and direct mono-exponential recovery of D_470_. In agreement with previous results, channel opening occurs at the UV/VIS silent transition P_390a_^M1^ to P_390b_^M2^ in two subsequent steps (8) that we can now attribute to different ion selectivities pinpointing to different pore conformations. Whereas during P_390b_^M2^ the short-lived O_1-early_ conducts almost exclusively protons, photocurrents of O_1-late_, that evolve upon reprotonation of the RSB and formation of P_520_^N^, are carried also by larger cations. Curiously, we observe a small positive charge displacement during P_520_^N^ at 0 mV and under symmetric ionic conditions (Fig. 2) that could result from an outward directed proton displacement following reprotonation of the retinal Schiff base. This might reflect the earlier proposed residual proton transfer in ChR2 (31) that was later associated with protonation changes of D156 (14). We note that the observed small charge transfer is barely visible after light-adaptation or any applied membrane potential. It occurs more than one order of magnitude later than the peak photocurrent in the proton pump bacteriorhodopsin (32) and differs from fast charge transfer observed in other channelrhodopsins (33).

In the second reaction path (forming a transition to the *syn*-cycle) illumination of D_470_ (C_1_) results in all-*trans*,C=N-*anti* → 13-*cis*,C=N-*syn* isomerization and direct formation of P_480_ (C_2_) which is also photo reactive. Branching of the photocycle has been considered before, to explain the bi-exponential decay of the conductive state P_520_ (14, 34, 35) and was assumed to involve an early all-*trans*,C=N-*anti* → 13-*cis*,C=N-*syn* isomerization based on NMR and low temperature Raman measurements (19). Using isotopically labelled retinals and vibrational spectroscopy, we proved that the early all-*trans*,C=N-*anti* → 13-*cis*,C=N-*syn* double isomerization (also at ambient temperatures) causes early formation of P_480_. Consequently, the slowly decaying P_480_ does not represent a late photocycle intermediate of the *anti*-cycle–as proposed in several previous publications (6, 14, 34)–but is the result of a reaction branching that occurs directly after excitation of D_470_ (C_1_), possibly already during the excited state lifetime. As shown by MD simulation, P_480_ features a pre-opening of the central gate allowing water influx, but remains non-conductive, because the inner gate is still closed as previously proposed for the E90R chloride-conducting mutant (20).

In a third reaction path (denoted as the *syn*-cycle), photoactivation of P_480_ (C_2_) initiates the *syn*-cycle. Here we identified P*_520_^N^ as the conductive state O_2_. Under continuous illumination, P*_520_^N^ accumulates due to its slow decay rate and significantly contributes to stationary photocurrent especially at high pH. Consequently P*_520_^N^ accumulation accounts for the bi-exponential channel-closing kinetics and the evolution of proton conductance during continuous illumination (4, 36). Comparing P*_520_^N^ accumulation in our FTIR measurements with the small photocurrent amplitude of O_2_ in our electrophysiological recordings indicates a significantly reduced conductance of O_2_ (P*_520_^N^) compared to O_1-early_ (P_390b_^M2^) and O_1-late_ (P_520_^N^) as previously predicted (9, 10). This explains now the photocurrent inactivation ChR2 during continuous illumination. It also explains the remarkably small shifts of the action spectra of the dark- (470 nm) and light-adapted (480 nm) protein (37). The reduced sodium selectivity of P*_520_^N^(O_2_) compared to P_520_^N^ (O_1-late_) indicates important differences in the open pore structures of both conducting states further supported by distinct FTIR spectra for P*_520_^N^ and P_520_^N^. In summary, the much slower decay of the *syn*-cycle with reduced conductance accumulates P_480_ and explains thereby now the inactivation

Finally, we hypothesize that E90 is crucial for proton conductance in both photocycles and constitutes the key determinant for ion selectivity changes during continuous illumination with an intriguing double function depending on its protonation state before and after light-adaptation. In the *anti*-cycle, E90 might be directly involved in proton transport as a proton shuttle or in the organization of water molecules that both bridge the distinct water-filled cavities seen in the dark state crystal structure of ChR2 (28). In this scenario E90 would favor proton selectivity, forming either a direct or indirect short-cut for protons that cannot be taken by larger cations at a similar efficiency. By similar means, the outer pore glutamates E139 and E143 in the highly proton-selective channelrhodopsin Chrimson (38) or D112 in voltage-gated proton channel Hv1 (39) were also shown to contribute to proton conductance and selectivity. Although the selectivity filter of ChR2 localized in the central gate might be more permissive for larger cations than that of Chrimson localized in the outer gate (40), in both cases substitution of essential glutamates (E90 in ChR2 and E139 in Chrimson) by equally titratable histidines preserved proton selectivity, whereas substitution with the non-titratable glutamine or alanine impaired proton conductance (13, 38). In the *anti*-cycle, channel opening occurs although E90 stays protonated for the entire gating process. Accordingly recent 4 µs MM-simulations on an ChR2 homology model based on the C1C2 chimera structure (PDB-ID: 3ug9 (29)) showed impressively that minor hydrogen bond changes in the central gate region were sufficient to promote water invasion in the same time range (28). We now observed a similar hydrogen bond rearrangement of E90 toward E123 in our P_390_ simulations for the ChR2 WT structure (Fig. S12) (27), that at the longer µs simulation times allowed water influx.

Once deprotonated E90 forms a salt bridge with the adjacent K93 and completely opens the central gate, promoting helix hydration from the extracellular site as shown in our MD simulations (Fig. 5). Although the proton acceptor of E90 has not been identified yet, the close proximity to water molecules as indicated by our MD-Simulations might allow fast proton diffusion into the bulk phase. Water molecules might serve as proton shuttles as they can become transiently protonated as previously shown for bacteriorhodosin (5). It is essential to note that pore hydration due to hydrogen bond changes of E90 in P_390_ differs from pore hydration due E90 deprotonation and salt-bridge formation with K93 that we proposed earlier in our E90-Helix-tilt model (15). Without considering parallel photocycles we initially attributed early pore hydration due to E90 deprotonation to a pre-gating step in a linear photocycle. At the same time early E90 deprotonation has been challenged by measurements of partly light-adapted ChR2 arguing that E90 is only deprotonated during the lifetime of P_480_ that was assumed to be a late intermediate of a linear photocycle or a late branching reaction (8, 14, 41). Comparing FTIR measurements of dark- and light-adapted ChR2 we were able to resolve the controversy regarding E90 deprotonation. P_480_ is formed in an ultrafast branching reaction that leads to the very fast deprotonation of E90. The splitting of the photocycle into the *anti-* and *syn*-branches forms the basis for the light-adaptation of ChR2. In the E90Q mutant all-*trans*,C=N-*anti* → 13-*cis*,C=N-*syn* isomerization still occurs but can no longer trigger deprotonation of residue 90. Consequently, in E90Q pore hydration in the *syn*-cycle is reduced and conformational changes during formation of P*_520_^N^ are no longer sufficient to support passive proton flux in the *syn*-cycle. As the *syn*-photocylce is still populated in the E90Q mutant but nonconductive, photocurrents still inactivate during continuous illumination at a degree determined by the relative rate constants. As an essential revision of our previous E90-Helix-tilt model, we therefore reassign early pore hydration to the light-dependent transition into the *syn*-cycle. Deprotonation of E90 and subsequent pore hydration do not constitute a pre-gating step for the highly conductive open channel intermediates O_1-early_ and O_1-late_ of the dark-adapted C=N-*anti*-cycle but instead prepare low proton conductance of P*_520_^N^ in the *syn-* cycle.

In a combined study of single-turnover electrophysiology and FTIR and Raman spectroscopy with isotopic retinal labelling, site-directed mutagenesis, and MD simulation, we developed a unifying two-photocycle model that simplifies and embraces previous kinetic models and completely resolves the channel gating, light-adaptation and temporal changes in ion selectivity. Identifying the corresponding molecular transitions, we may facilitate future protein engineering of channelrhodopsin variants with reduced or improved photocurrent inactivation for optogenetic applications, requiring either stable response to continuous illumination or a transient response to light switching. Early photocycle branching by alternative retinal isomerization and the corresponding large conformational protein changes that do not directly lead to channel opening will need careful consideration for the interpretation of molecular gating transitions observed in time resolved spectroscopy and crystallography.

## Materials and methods

Yeast culture: *Pichia pastoris* strain SMD1163 cells (kindly gifted by C. Bamann) containing the pPIC9KChR2His10 construct were precultured in BMGY medium (42). Expression of ChR2 was induced in BMMY medium containing 2.5 µM all-*trans* retinal (either ^12^C, ^13^C_14_,^13^C_15_-labelled or ^13^C_10_,^13^C_11_-labelled) and 0.00004% biotin at an initial OD_600_ of 1 and at 30 °C and 120 rpm. Cells were harvested at an OD_600_ of 20 by centrifugation.

Membrane preparation and protein purification: Cells were disrupted using a BeadBeater (Biospec Products) and membranes were isolated by ultracentrifugation. Homogenized membranes were solubilized with 1% decyl maltoside overnight. ChR2 purification was done by Ni-NTA affinity chromatography and subsequent gelfiltration using a HiLoad 16/600 Superdex 200 pg column (GE).

### Reconstitution of ChR2 into DPPC or EggPC

The purified ChR2 was reconstituted into 1,2-dipalmitoyl-*sn*-glycero-3-phosphocholine (DPPC, Avanti Polar Lipids) or EggPC (Avanti Polar Lipids). The lipids were solubilized with 2% cholate in 20 mM Hepes pH 7.5, 100 mM NaCl and 1 mM MgCL_2_ by incubation at 50 °C for 10 min. Solubilized lipids and purified ChR2 were mixed at a 2:1 ratio (lipid:protein, w:w) and incubated for 20 min. Detergent was removed overnight either by adsorption on Bio-Beads SM 2 (BioRad) or by dialysis.

The resulting suspension containing proteoliposomes and buffer was then ultracentrifuged at 200000 g for 2 h and the pellet was then transferred and squeezed between two CaF_2_-slides to obtain an optical path length between 5–10 µm. This sample was then placed in a vacuum tight cuvette.

### Preparation of HEK cells

Electrophysiological recordings were performed on stably expressing *ChR2*-mVenus fusion construct HEK cell line (31) as previously described in detail (43). Briefly, HEK cells were cultured at 5% CO_2_ and 37 °C in Dulbecco’s minimal essential medium supplemented with 10% fetal bovine serum, 100 µg/mL penicillin/streptomycin (Biochrom, Berlin, Germany), 200 µg/mL zeocin, and 50 µg/mL blasticidin (Thermo Fischer Scientific, Waltham, MA). Cells were seeded onto poly-lysine coated glass coverslips at a concentration of 1 × 10^5^ cells/mL and supplemented with a final concentration of 1 μM all-trans retinal (Sigma-Aldrich, Munich, Germany). Induction of *ChR2*-mVenus expression was induced by addition of 0.1 µM tetracyclin (Thermo Fischer Scientific).

### Patch-clamp experiments in HEK293 cells

Patch pipettes were pulled using a P1000 micropipette puller (Sutter Instruments, Novato, CA), and fire-polished. Pipette resistance was 1.5 to 2.5 MΩ. A 140 mM NaCl agar bridge served as reference (bath) electrode. In whole-cell recordings, membrane resistance was typically > 1 GΩ, while access resistance was below 10 MΩ. Pipette capacity, series resistance, and cell capacity compensation were applied. All experiments were carried out at 23 °C. Signals were amplified (AxoPatch200B), digitized (DigiData1400), and acquired using Clampex 10.4 Software (all from Molecular Devices, Sunnyvale, CA). Holding potentials were varied in 15 mV steps between −60 and +30 mV. Extracellular buffer exchange was performed manually by adding at least 5 mL of the respective buffer to the recording chamber (500 µL chamber volume) while a Ringer Bath Handler MPCU (Lorenz Messgerätebau, Katlenburg-Lindau, Germany) maintained a constant bath level. Standard bath solutions contained 110 mM NaCl, 1 mM KCl, 1 mM CsCl, 2 mM CaCl_2_, 2 mM MgCl_2_, and 10 mM HEPES at pH_e_ 7.2 (with glucose added up to 310 mOsm). Standard pipette solutions contained 110 mM NaCl, 1 mM KCl, 1 mM CsCl, 2 mM CaCl_2_, 2 mM MgCl_2_, 10 mM EGTA, and 10 mM HEPES at pH_i_ 7.2 or 10 mM TRIS at pH_i_ 9.0 (glucose was added up to 290 mOsm). For ion selectivity measurements, either NaCl was replaced by 110 mM NMDGCl, or extracellular pH was adjusted to pH_e_ 9.0 by buffering with 10 mM TRIS instead of HEPES.

Continuous light was generated using Polychrome V light source (TILL Photonics, Planegg, Germany) set to 470 ± 7 nm. Light exposure was controlled with a programmable shutter system (VS25 and VCM-D1, Vincent Associates, Rochester, NY). The Polychrome V light intensity was 3.4 mW/mm^2^ in the sample plane, measured with a calibrated optometer (P9710, Gigahertz Optik, Türkenfeld, Germany). Light intensities were calculated for the illuminated field of the W Plan-Apochromat 40x/1.0 DIC objective (0.066 mm^2^, Carl Zeiss, Jena, Germany). For delivery of 470 nm ns-laser pulses, an Opolette polette HENd:YAG laser/OPO system (OPOTEK, Carlsbad, CA) was coupled/decoupled into a M37L02 multimode fiber patch cable with a modified KT110/M free-space-to-fiber coupler using AC127 019 A ML achromatic doublets (Thorlabs, Newton, NJ). Single pulses were selected using a LS6ZM2 shutter (Vincent Associates). Laser intensity was set to 5% using the built-in motorized variable attenuator, resulting in a pulse energy of 100 ± 20 µJ/mm^2^. Pulse energies were measured with a calibrated S470C thermal power sensor and a PM100D power and energy meter (Thorlabs) after passing through all the optics. Actinic light was coupled into an Axiovert 100 microscope (Carl Zeiss) and delivered to the sample using a 90/10 beamsplitter (Chroma, Bellows Falls, VT). To toggle between activation with the laser and the Polychrome V, a BB1 E02 broadband dielectric mirror mounted on a MFF101/M motorized filter flip mount (Thorlabs) was used. Data was filtered at 100 kHz and sampled at 250 kHz. Due to minimal timing uncertainties, each acquired sweep was time shifted post measurements to align it with the rising edges of the Q-switch signals of the activating laser pulses. Photocurrents were binned to 50 logarithmically spaced data points per temporal decade with custom-written Matlab script (MathWorks, Natick, MA).

### FTIR-Experiments

To gain insight into the changes upon illumination, we performed time resolved FTIR difference spectroscopy at 15 °C. For the continuous light experiments, the sample was illuminated with a blue LED (λ_max_ = 465 nm) for 5 s. Spectra were recorded before switching on the light (reference), during illumination (accumulation of P_Stat_ for 5 s) and after switching off the light (decay of P_Stat_ for 500 s) using the conventional rapid scan mode of the spectrometer. Difference spectra were calculated using the Beer-Lambert law which results in positive photo product bands and negative educt bands in the difference spectra. For the single-turnover measurements, the sample was illuminated with a short laser pulse of an excimer laser-driven dye laser (dye: Coumarin 102, λ_max_: 475 nm, pulse width: approx. 50 ns). Conventional rapid scan experiments (time resolution: approx. 10 ms, spectral resolution: 4 cm^−1^) were performed with a sufficient relaxation time (*t*_relax_ = 200 s, f_flash_ = 0.005 Hz) between the flashes to allow the D_470_ to significantly repopulate ([D_470_] became approx. 96%).

For a comparison with the photocycle under the “shortcut condition” the flash frequency was increased (*t*_relax_ = 5 s, *f*_flash_ = 0.2 Hz). Using this approach, equilibrium between D_470_ and P_480_ emerges and the ChR2 molecules start the photocycle from both states. The datasets were then analyzed by a global fitting routine as presented previously (13, 15, 44, 45) to isolate the decay associated amplitude spectra of the transitions involved in D_470_ recovery. To get access to the earlier Intermediates of the dark-adapted ([D_470_] approx. 91%) photocycle of H134R (Fig. 3A), step-scan measurements were performed with a light pulse repetition rate of 0.007 Hz (*t*_relax_ = 140 s detector rise time: 50 ns, resolution: 8 cm^−1^, wavenumber range: 0–1974 cm^−1^) as already published for the WT (15). One Measurement was completed after 22 h, and approx. 15 measurements were averaged to give the final result. H134R used for the step scan was expressed in COS-cells and prepared as described in our earlier publication (15).

### Raman Experiments

The Raman experiments were performed with samples that were prepared exactly as the ones for the rapid-scan FTIR-experiments, but with a higher optical path length (20–50 µm). The room temperature was approx. 18 °C. We used the Raman microscope XPloRA One (HORIBA Scientific, 3880 Park Avenue, Edison, New Jersey) to scan the sample. In order to prevent sample degradation due to long illumination, we performed measurements at a 785-nm excitation wavelength to ensure the lowest possible photoexcitation of the sample and a sufficient enhancement of the Raman signal due to the pre-resonant Raman effect. Laser power at the sample position was 28 mW. A 50× objective (Olympus LCPLN-IR) was used, resulting in a confocal volume of approx. 1 µm^3^ in the sample plane.

To excite the D_470_ state of the sample and create a photoproduct with a high P_480_ fraction, the sample was illuminated with an external blue light source (100 W halogen lamp filtered with a 470 nm filter coupled into the observation beam path of the microscope). Sample illumination was controlled by a shutter between lamp and sample.

To ensure that the measured photoproduct spectra are free of contamination by their preceding intermediates (P_520_^N^ and P_390_^M^) under continuous illumination, a controlled illumination/relaxation experiment was performed:

Laser On+illumination on (Formation of P_Stat_)

Wait 0.5 s

Acquisition of spectrum (*t*_Int_ = 2 s). P_Stat_ is measured.

Laser On+illumination off (slow relaxation of P_Stat_: *t*_1/2_ approx. 40 s)

Wait 0.5 s

Acquisition of spectrum (*t*_Int_ = 2 s). High fraction of P_480_ free of illumination artifacts is measured (P’_Stat_).

Go to next position on the sample

This procedure was repeated for a 15 × 15 spot matrix (pixel spacing: 5 µm) of the sample and the acquired spectra were averaged for each illumination condition.

Next, the illumination was stopped and a relaxation phase of at least 5 minutes was commenced to allow full relaxation of the generated P_480_. The dark state D_470_ was measured for the same spot matrix (*t*_Int_ = 2 s).

To obtain the pure lamp artifact, the same area was measured without the laser with only illumination of the sample. This spectrum was later subtracted from the spectra measured under continuous illumination.

The complete protocol was performed for three spectral regions (center wavenumbers: 900 cm^−1^, 1250 cm^−1^, and 1550 cm^−1^) which were then combined to obtain the complete spectrum ranging from 650 cm^−1^ to 1700 cm^−1^ with a wavenumber spacing of approx. 0.2 cm^−1^ and a nominal resolution of 0.6 cm^−1^.

### MD simulations

The MD simulations were performed according to our previous reports (13, 15) except for the force field and the GROMACS version used. We used the OPLS/AA all atom force field and GROMACS version 2016.3. A series of 5 × 100 ns independent and unrestrained MD simulations was performed for each protonation state of E90 with the respective chromophore configuration. The MD simulations were performed consecutively using the resulting structures of dark-adapted ChR2 (all-*trans*,C=N-*anti* with protonated E90) given by MD simulations as a starting point for the isomerization (see below). Each MD simulation was initiated using a different temperature seed number to generate the random distribution of starting velocities.

### Water dynamic and run-average structure

The water dynamic and run-average structures were performed according to our previous report (15).

### Retinal isomerization

Retinal all-*trans*,C=N-*anti* to 13-*cis*,C=N-*anti* isomerization was performed as described earlier (15), achieved via by the following scheme. The torsion angles of the C13=C14 and C=N double bonds were tilted counterclockwise in 20° steps starting in the range 0–180°. For each tilting step, the Retinal + K257 (without backbone) atoms were maintained as a freeze group and the rest of the simulation system was allowed to relax in a 10-ns unrestrained MD-simulation as described above.

The resulting 13-*cis* retinal structures served as starting structures for the MD simulations of the different intermediates (P_500_^K^, P_480_ and P_480_-E90p) with protonated and deprotonated E90. For the starting structures for the P_390a_^M1^ intermediate, we used the final structures of the P_500_^K^ simulations after Schiff base deprotonation and D253 protonation.

## Acknowledgments

We thank Harald Chrongiewski and Gabi Smuda for technical assistance. We also thank Mathias Lübben and Till Rudack for helpful discussions. This work was supportet by the Deutsche Forschungsgemeinschaft DFG Priority Programme SPP 1926.

We thank Maila Reh, Altina Klein and Tharsana Tharmalingam for technical assistance. We also thank Christiane Grimm, Joel Kaufmann and Franz Bartl for fruitful discussions. This work was supported by the Deutsche Forschungsgemeinschaft DFG [SFB1078 (B2) and the Cluster of Excellence Unifying Concepts in Catalysis, UniCat, BIG-NSE (Johannes Vierock) and E4 (Peter Hegemann)]. Peter Hegemann is Senior Research Professor of the Hertie Foundation.

## Author contributions

Jens Kuhne and Johannes Vierock contributed equally and either has the right to list himself first in bibliographic documents. The study was designed by Jens Kuhne, Johannes Vierock, Peter Hegemann and Klaus Gerwert. Protein expression and purification was established by Konstantin Gavriljuk und carried out by Max-Aylmer Dreier. The setup for time-resolved FTIR experiments was developed and established by Jens Kuhne. The setup for Raman experiments was developed and established by Dennis Peterson and measurements were carried out by Jens Kuhne. Raman results were analyzed with the help of Samir F. El-Mashtoly. The biomolecular simulations were performed and analyzed by Stefan Alexander Tennigkeit. The setup for single-turnover electrophysiological experiments was developed and established by Jonas Wietek. and Johannes Vierock. The electrophysiological experiments were performed and analyzed by Johannes Vierock.

Jens Kuhne, Johannes Vierock, Stefan Tennigkeit, Peter Hegemann an Klaus Gerwert wrote the manuscript with the help of Max-Aylmer Dreier and Jonas Wietek.

## References

1. Zhang F, et al. (2011) The Microbial Opsin Family of Optogenetic Tools. Cell 147(7):1446–1457.

2. Scheib U, et al. (2015) The rhodopsin-guanylyl cyclase of the aquatic fungus Blastocladiella emersonii enables fast optical control of cGMP signaling. Sci Signal 8(389):rs8.

3. Boyden ES, Zhang F, Bamberg E, Nagel G, Deisseroth K (2005) Millisecond-timescale, genetically targeted optical control of neural activity. Nat Neurosci 8(9):1263–1268.

4. Nagel G, et al. (2003) Channelrhodopsin-2, a directly light-gated cation-selective membrane channel. Proc Natl Acad Sci 100(24):13940–13945.

5. Garczarek F, Gerwert K (2005) Functional waters in intraprotein proton transfer monitored by FTIR difference spectroscopy. Nature 439(7072):109–112.

6. Ritter E, Stehfest K, Berndt A, Hegemann P, Bartl FJ (2008) Monitoring Light-induced Structural Changes of Channelrhodopsin-2 by UV-visible and Fourier Transform Infrared Spectroscopy. J Biol Chem 283(50):35033–35041.

7. Bamann C, Kirsch T, Nagel G, Bamberg E (2008) Spectral characteristics of the photocycle of channelrhodopsin-2 and its implication for channel function. J Mol Biol 375(3):686–694.

8. Lórenz-Fonfría VA, et al. (2015) Temporal evolution of helix hydration in a light-gated ion channel correlates with ion conductance. Proc Natl Acad Sci U S A 112(43):E5796-5804.

9. Hegemann P, Ehlenbeck S, Gradmann D (2005) Multiple photocycles of channelrhodopsin. Biophys J 89(6):3911–3918.

10. Nikolic K, et al. (2009) Photocycles of Channelrhodopsin-2. Photochem Photobiol 85(1):400–411.

11. Valentini A, et al. (2017) Optomechanical Control of Quantum Yield in Trans–Cis Ultrafast Photoisomerization of a Retinal Chromophore Model. Angew Chem Int Ed 56(14):3842–3846.

12. Verhoefen M-K, et al. (2010) The Photocycle of Channelrhodopsin-2: Ultrafast Reaction Dynamics and Subsequent Reaction Steps. ChemPhysChem 11(14):3113–3122.

13. Eisenhauer K, et al. (2012) In channelrhodopsin-2 Glu-90 is crucial for ion selectivity and is deprotonated during the photocycle. J Biol Chem 287(9):6904–6911.

14. Lórenz-Fonfría VA, et al. (2013) Transient protonation changes in channelrhodopsin-2 and their relevance to channel gating. Proc Natl Acad Sci:201219502.

15. Kuhne J, et al. (2015) Early Formation of the Ion-Conducting Pore in Channelrhodopsin-2. Angew Chem Int Ed 54(16):4953–4957.

16. Gerwert K, Freier E, Wolf S (2014) The role of protein-bound water molecules in microbial rhodopsins. Biochim Biophys Acta 1837(5):606–613.

17. Ritter E, Piwowarski P, Hegemann P, Bartl FJ (2013) Light-dark adaptation of channelrhodopsin C128T mutant. J Biol Chem 288(15):10451–10458.

18. Becker-Baldus J, et al. (2015) Enlightening the photoactive site of channelrhodopsin-2 by DNP-enhanced solid-state NMR spectroscopy. Proc Natl Acad Sci 112(32):9896–9901.

19. Bruun S, et al. (2015) Light–Dark Adaptation of Channelrhodopsin Involves Photoconversion between the all-*trans* and 13-*cis* Retinal Isomers. Biochemistry 54(35):5389–5400.

20. Wietek J, et al. (2014) Conversion of channelrhodopsin into a light-gated chloride channel. Science 344(6182):409–412.

21. Kaufmann JCD, et al. (2017) Proton transfer reactions in the red light-activatable channelrhodopsin variant ReaChR and their relevance for its function. J Biol Chem 292(34):14205–14216.

22. Berndt A, et al. (2011) High-efficiency channelrhodopsins for fast neuronal stimulation at low light levels. Proc Natl Acad Sci 108(18):7595–7600.

23. Gerwert K, Siebert F (1986) Evidence for light-induced 13-cis, 14-s-cis isomerization in bacteriorhodopsin obtained by FTIR difference spectroscopy using isotopically labelled retinals. EMBO J 5(4):805–811.

24. Smith SO, et al. (1984) Determination of retinal Schiff base configuration in bacteriorhodopsin. Proc Natl Acad Sci 81(7):2055–2059.

25. Babitzki G, Mathias G, Tavan P (2009) The Infrared Spectra of the Retinal Chromophore in Bacteriorhodopsin Calculated by a DFT/MM Approach. J Phys Chem B 113(30):10496–10508.

26. Lórenz-Fonfría VA, Muders V, Schlesinger R, Heberle J(2014) Changes in the hydrogen-bonding strength of internal water molecules and cysteine residues in the conductive state of channelrhodopsin-1. J Chem Phys 141(22):22D507.

27. Volkov O, et al. (2017) Structural insights into ion conduction by channelrhodopsin 2. Science 358(6366):eaan8862.

28. Ardevol A, Hummer G (2018) Retinal isomerization and water-pore formation in channelrhodopsin-2. Proc Natl Acad Sci 115(14):3557–3562.

29. Kato HE, et al. (2012) Crystal structure of the channelrhodopsin light-gated cation channel. Nature 482(7385):369–374.

30. Ruffert K, et al. (2011) Glutamate residue 90 in the predicted transmembrane domain 2 is crucial for cation flux through channelrhodopsin 2. Biochem Biophys Res Commun 410(4):737–743.

31. Feldbauer K, et al. (2009) Channelrhodopsin-2 is a leaky proton pump. Proc Natl Acad Sci 106(30):12317–12322.

32. Keszthelyi L., Ormos P. (2001) Electric signals associated with the photocycle of bacteriorhodopsin. FEBS Lett 109(2):189–193.

33. Sineshchekov OA, Govorunova EG, Wang J, Li H, Spudich JL (2013) Intramolecular Proton Transfer in Channelrhodopsins. Biophys J 104(4):807–817.

34. Bamann C, Gueta R, Kleinlogel S, Nagel G, Bamberg E (2010) Structural guidance of the photocycle of channelrhodopsin-2 by an interhelical hydrogen bond. Biochemistry 49(2):267–278.

35. Stehfest K, Hegemann P (2010) Evolution of the Channelrhodopsin Photocycle Model. ChemPhysChem 11(6):1120–1126.

36. Berndt A, Prigge M, Gradmann D, Hegemann P (2010) Two Open States with Progressive Proton Selectivities in the Branched Channelrhodopsin-2 Photocycle. Biophys J 98(5):753–761.

37. Lin JY, Lin MZ, Steinbach P, Tsien RY (2009) Characterization of engineered channelrhodopsin variants with improved properties and kinetics. Biophys J 96(5):1803–1814.

38. Vierock J, Grimm C, Nitzan N, Hegemann P (2017) Molecular determinants of proton selectivity and gating in the red-light activated channelrhodopsin Chrimson. Sci Rep 7(1):9928.

39. Dudev T, et al. (2015) Selectivity Mechanism of the Voltage-gated Proton Channel, HV1. Sci Rep 5:10320.

40. Oda K, et al. (2018) Crystal structure of the red light-activated channelrhodopsin Chrimson. Nat Commun 9(1):3949.

41. Saita M, et al. (2018) Photoexcitation of the P4480 State Induces a Secondary Photocycle That Potentially Desensitizes Channelrhodopsin-2. J Am Chem Soc 140(31):9899–9903.

42. Radu I, et al. (2009) Conformational Changes of Channelrhodopsin-2. J Am Chem Soc 131(21):7313–7319.

43. Grimm C, Vierock J, Hegemann P, Wietek J (2017) Whole-cell Patch-clamp Recordings for Electrophysiological Determination of Ion Selectivity in Channelrhodopsins. JoVE J Vis Exp (123):e55497–e55497.

44. Müller K-H, Plesser T (1991) Variance reduction by simultaneous multi-exponential analysis of data sets from different experiments. Eur Biophys J 19(5):231–240.

45. Hessling B, Souvignier G, Gerwert K (1993) A model-independent approach to assigning bacteriorhodopsin’s intramolecular reactions to photocycle intermediates. Biophys J 65(5):1929–1941.

