## Supplementary Information for "A unifying photocycle model for light adaptation and temporal evolution of cation conductance in Channelrhodopsin-2"

**This file includes:**

Supplementary Note 1 to 7
Figs. S1 to S14
Tables S1
References for SI reference citations

### Supplementary Information Text

**Supplementary Note 1: HPLC retinal extraction experiments.** Previously published HPLC retinal extraction experiments showed only minor changes in the extracted retinal isomers after light adaptation (1). In contrast our FTIR and Raman data (Fig. 3-4 in the main text and Fig. S2-S6) indicate large structural changes of the retinal upon light adaptation.

To resolve this discrepancy between vibrational spectroscopy and HPLC results, we performed HPLC retinal extraction experiments on solubilized ChR2 as well as on lipid reconstituted ChR2 (see Table S1). For dark adapted solubilized samples the extraction contains approx. 74 % all-*trans* retinal, 25 % 13-*cis* retinal and a minor fraction of 9-*cis* and 11-*cis* species. Upon illumination, these values are only slightly altered. This agrees well with the approximate 70:30 ratio of all-*trans* to 13-*cis* retinal for dark adapted protein that was found in a previous study (1). If the sample is reconstituted in EggPC an approx. 79 % /19 % all-*trans*/13-*cis* mixture is found in the extraction of the dark adapted samples. Upon illumination a loss of over 20 percentage points of the all-trans retinal is visible. The 13-*cis* species component increases about 10 percentage points and the values of presumably 11-*cis* and 9-*cis* retinals are slightly increased. The change of the isomeric composition of the extraction upon ChR2 light adaptation is less intense if DPPC is used as a lipid for reconstitution.

Obviously, the environment influences the results of the HPLC retinal extraction experiments. Light adaptation seems to be impeded in solubilized samples which might be the reason for the discrepancy between vibrational spectroscopy and HPLC measurements.

Further the values found for the dark adapted sample strongly differ from NMR results of dark adapted ChR2 reconstituted in lipids (2, 3). Since NMR detects the retinal conformation inside the sample, whereas the HPLC data is only available for the extracted retinal, NMR data has to be regarded as more precise. The harsh conditions applied to extract the retinal out of the sample might alter the result drastically (3).

**Supplementary Note 2: E90 deprotonation triggers water influx.** We compared $P_{480}$-minus-$D_{470}$ FTIR difference spectra of the Wild type and variants that prevent E90 deprotonation (See Fig. S1). We found that in addition to the missing or less pronounced E90 bands (carbonyl stretch at 1718 $cm^{-1}$ and asymmetric/symmetric carboxylate stretches at approx. 1514/1381 $cm^{-1}$) the helix hydration marker bands at 1660/1650 $cm^{-1}$ is reduced. Water influx upon $P_{480}$ formation seems to be coupled to E90 deprotonation.

**Supplementary Note 3: Monitoring $D_{470}$ and $P_{480}$ using FTIR and Raman Spectroscopy.** In order to assign the retinal bands of ChR2, we performed FTIR difference spectroscopic measurements which were complemented by pre-resonance Raman spectra. Upon continuous blue actinic light illumination of the $D_{470}$ state, the photo stationary state $P_{Stat}$ is formed(4). The $P_{Stat}$-minus-$D_{470}$ FTIR difference spectrum (orange spectrum) shown in Fig. S2A exhibits strong positive and negative bands that correspond to $P_{Stat}$ and $D_{470}$, respectively. Two seconds after switching off the light the shape of the difference spectrum is only slightly altered ($P'_{Stat}$-minus-$D_{470}$, red spectrum). A global fit analysis of the $P_{Stat}$ decay (Fig. S2D) reveals a small fraction of a fast process ($t_{1/2}$=250 ms) representing most likely channel closing and a main fraction with a half-life of 40 s which is the $P_{480}$ decay(5, 6). This means that the $P'_{Stat}$-minus-$D_{470}$ difference spectrum measured 2 seconds after the end of illumination (red spectrum in Fig. S2A) is exactly a $P_{480}$-minus-$D_{470}$ difference spectrum.

In addition we performed pre-resonance Raman experiments on FTIR-samples (with higher optical pathlength) to maintain similar experimental conditions. A 785 nm excitation was used in order to generate a pre-resonance condition without producing photo-cycle intermediates. In Fig. S2B the Raman spectrum of $D_{470}$ (blue spectrum) is shown. In contrast to the aforementioned study (1), considerable spectral changes were observed when additional blue light (λ=470 nm) was applied on the sample to accumulate $P_{Stat}$ (orange spectrum). The intensity of Raman bands at 1273 $cm^{-1}$ and 1157 $cm^{-1}$ decreases significantly and a new band at 1179 $cm^{-1}$ appears upon illumination. The magnitude of the spectral changes depends on the illumination intensity; however saturation was detected already at moderate brightness of the external light source (Fig. S2B inset). The spectral signature of $P_{Stat}$ is only slightly altered 2 s after the light is switched off (P'$_{Stat}$, red spectrum). The Raman difference spectra of $P_{Stat}$-minus-P'$_{Stat}$ and P'$_{Stat}$-minus-$D_{470}$ (Fig. S2C) match well with that measured by FTIR as well as the decay associated amplitude spectra (Fig. S2C) showing that the same processes (open state and $P_{480}$-decay) can be monitored using both techniques.

The negative band of E90 at 1718 $cm^{-1}$ has been assigned to the protonated state of E90 in $D_{470}$ (4, 6). It was shown that in $P_{480}$, E90 is in the deprotated state, since no corresponding positive carbonyl band has been found(5). The negative band at 1662 $cm^{-1}$ in the amide region is partly composed of C=O oscillators of dehydrated transmembrane helices in $D_{470}$ (7). Further the N-H- stretching vibration of the protonated SB at 1658 $cm^{-1}$ contributes to this intense $D_{470}$ band (2). The positive $P_{480}$ band at 1650 $cm^{-1}$ has been assigned to C=O oscillators of hydrated transmembrane helices, indicating water influx into the protein (7). The C=C-stretching bands of the retinal polyene chain are expected 1500 $cm^{-1}$ and 1600 $cm^{-1}$. In this region a negative $D_{470}$ band at 1547 $cm^{-1}$ has been detected. The corresponding positive C=C-stretch band of $P_{480}$ is located at 1560 $cm^{-1}$ which is unusual since the absorption maximum is red shifted in $P_{480}$ (8). In the conformation-dependent fingerprint region strong C-C-stretching bands of the retinal polyene chain are visible. Four negative bands at 1244 $cm^{-1}$, 1234 $cm^{-1}$, 1202 $cm^{-1}$, 1186 $cm^{-1}$, and two positive bands at 1176 $cm^{-1}$ and 1154 $cm^{-1}$ indicate that the conformation of the retinal is strongly altered in the $D_{470}$ to $P_{480}$ transition. In contrast, minor changes of the chromophore were detected in pre-resonance Raman measurements during the formation of $P_{Stat}$ (1).

The bands at 1273 $cm^{-1}$, 1244 $cm^{-1}$ and 1234 $cm^{-1}$ are negative in the P'$_{Stat}$-minus-$D_{470}$ Raman and FTIR difference spectra. Therefore these bands are marker bands for the $D_{470}$ state that is obviously incompletely depopulated upon continuous illumination which can be better seen by the residual amount of the 1273 $cm^{-1}$ band in the P'$_{Stat}$ spectrum. Since the intense positive band at 1179 $cm^{-1}$ is visible in both the P'$_{Stat}$-minus-$D_{470}$ (Raman) and the $P_{480}$-minus-$D_{470}$ (FTIR) difference spectra it is considered as a Raman- and FTIR-marker band for $P_{480}$. The spectrum of P'$_{Stat}$ contains approximately similar amounts of $P_{480}$ and $D_{470}$ which cannot be seen in difference spectra due to the subtraction of absolute spectra. The residual amount of $D_{470}$ can be removed by subtraction of a scaled pure $D_{470}$ spectrum from the spectrum of P'$_{Stat}$ using the $D_{470}$-marker band at 1273 $cm^{-1}$ as a guide. Fig. S4 and main text Fig. 4 show the resulting pure spectra of $P_{480}$ (red) and $D_{470}$ (black). Since the retinal bands are selectively enhanced due to the pre-resonance Raman Effect, most of the visible bands can be attributed to the retinal. The Raman band assignments will be discussed in Fig. S6. The overall band patterns agree surprisingly well with the published spectra of the 13-*cis*,C=N-*syn*- and all-*trans*,C=N-*anti* components of the $D_{app}$ state that is formed within 30 µs after the onset of illumination (2).

Similar spectra can be generated by applying Non Negative Matrix Factorization (NNMF (9)) on a dataset measured under various illumination intensities. This procedure takes into account, that the Raman spectra are absolute spectra that are always positive. If two components are used, the new base spectra shown in Fig. S5A match well the ones generated by the subtraction procedure (Fig. S4). The concentration profiles in Fig. S5B show nicely that only $D_{470}$ exists without additional illumination. The fraction of $P_{480}$ is generated upon illumination but saturates at max. 60 % even if the illumination intensity is further increased.

**Supplementary Note 4: Band assignment of the $P_{480}$ and $D_{470}$ bands.** In Fig. S6 the $P_{480}$ and $D_{470}$ bands in the Raman spectra and the $P_{480}$-minus-$D_{470}$ FTIR difference spectra are compared for unlabeled (black) and $^{13}C_{14}$-$^{13}C_{15}$-labelled (red) retinal. Since the sample with $^{13}C_{10}$-$^{13}C_{11}$-labelled retinal was only poorly reactive after reconstitution, only the $P_{480}$-minus-$D_{470}$ difference spectra are shown. However the labelling efficiency can be estimated as higher than 90% as indicated by the complete downshift of the negative band at 1273 $cm^{-1}$ upon $^{13}C_{10}$-$^{13}C_{11}$-labelling.

<u>C=N stretching region (1700 $cm^{-1}$- 1600 $cm^{-1}$):</u> The C=N-stretching band of $D_{470}$ is found at 1659 $cm^{-1}$ and can be clearly assigned to retinal due to its 16 $cm^{-1}$ downshift upon $^{13}C_{14}$-$^{13}C_{15}$-labelling. In $P_{480}$, that the C=N band broadens and is downshifted to 1630 $cm^{-1}$. Due to the loss in Raman intensity upon labeling a clear assignment of the shifted band is not possible. However it is tentatively assigned to 1610 $cm^{-1}$ in the FTIR spectrum because of the increased absorbance in the spectrum of the labeled sample.

<u>C=C stretching region (1500 $cm^{-1}$- 1600 $cm^{-1}$)</u>: The C=C-stretching band of $D_{470}$ at 1552 $cm^{-1}$ splits in the $^{13}C_{14}$-$^{13}C_{15}$-labelled sample. Such splitting upon $^{13}C_{14}$-$^{13}C_{15}$-labelling is also observed in crystals of all-trans retinal with PSB (10). In $P_{480}$ the C=C-stretching band of the unlabeled sample is slightly blue shifted to 1556 $cm^{-1}$. Upon labeling a 2 $cm^{-1}$ downshift of the band is observed.

<u>Fingerprint region (Combination of single bond ethylenic stretches, 1300 $cm^{-1}$ – 1100 $cm^{-1}$):</u> The marker band of $D_{470}$ at 1273 $cm^{-1}$ is most likely caused by the H11 in-plane rocking vibration coupled with the C-C-stretches of the retinal since it is sensitive to $^{13}C$ labeling at carbon positions $C_{10}$-$C_{11}$ (4 $cm^{-1}$ downshift) but not at $C_{13}$-$C_{14}$ (see Raman $P_{480}$-minus-$D_{470}$ difference spectrum in Fig. S6D). This band assignment is in agreement with published data of pure all-*trans* retinal PSB (10) and light adapted BR (11) where a band at identical position was found.

The $D_{470}$ bands at 1244 $cm^{-1}$ and 1235 $cm^{-1}$ are most likely due to delocalized anti-symmetric C-C modes of the chromophore in the all-*trans*,C=N-*anti* form. In $BR_{568}$ a band at 1254 $cm^{-1}$ has been assigned to the anti-symmetric combination of the $C_{12}$-$C_{13}$ + $C_{14}$-$C_{15}$-stretching and the $H_{14}$ rocking vibration. Since the $D_{470}$-band at 1234 $cm^{-1}$ is slightly downshifted upon $^{13}C_{14}$-$^{13}C_{15}$-labelling it is assigned to this combination mode in ChR2. These bands vanish in the $P_{480}$ spectra.

In light adapted bR the band at 1201 $cm^{-1}$ reflects the $C_{14}$-$C_{15}$ stretch vibration of the all-trans, C=N-*anti* retinal (11). In contrast this assignment is not valid for ChR2. Neither in the $D_{470}$ nor the $P_{480}$ spectrum the band at 1202 $cm^{-1}$ is significantly downshifted upon $^{13}C_{14}$-$^{13}C_{15}$-labelling. Instead one part of the $D_{470}$-band at 1186 $cm^{-1}$ is downshifted to 1169 $cm^{-1}$ as visible in FTIR and Raman spectra. This agrees well with the 19 $cm^{-1}$ downshift of the 1191 $cm^{-1}$ band as observed in crystals of all-*trans* retinal PSB (10). In contrast no such downshift is observed for the 1184 $cm^{-1}$ band of $P_{480}$. The $P_{480}$-band at 1179 $cm^{-1}$ is hardly affected by the $^{13}C_{14}$-$^{13}C_{15}$-labelling. But upon $^{13}C_{10}$-$^{13}C_{11}$-labelling a significant downshift of this band is observed in the FTIR and Raman $P_{480}$-minus-$D_{470}$ difference spectra. Therefore the $P_{480}$-band at 1179 $cm^{-1}$ is assigned to the $C_{10}$-$C_{11}$ stretch vibration. The $D_{470}$ band at 1157 $cm^{-1}$ shows a strange behavior. It vanishes upon $^{13}C_{14}$-$^{13}C_{15}$-labelling but is significantly downshifted if carbons $C_{10}$ and $C_{11}$ are labeled (see the Raman $P_{480}$-minus-$D_{470}$ difference spectrum). This has been also observed for the 1159 $cm^{-1}$ band of crystals of all-trans retinal with a PSB (10). Further no negative band in the FTIR difference spectrum is visible at 1159 $cm^{-1}$ as well as at the respective positions of the labeled sample. We therefore conclude that this band is caused by a solely Raman active mode that is dominated by the $C_{10}$-$C_{11}$ stretching vibration.

The positive $P_{480}$ band in the $P_{480}$-minus-$D_{470}$ FTIR difference spectra at 1154 $cm^{-1}$ is assigned to the $C_{14}$-$C_{15}$ stretching vibration of $P_{480}$ since it is about 14 $cm^{-1}$ downshifted upon $^{13}C_{14}$-$^{13}C_{15}$-labelling. This downshift is also visible in the $P_{480}$ Raman spectra although the Raman intensity is very low. Due to the overlap with the strongly Raman active $D_{470}$ band at 1157 $cm^{-1}$ this band is not visible in the Raman $P_{480}$-minus-$D_{470}$ difference spectrum. In $BR_L$ the $C_{14}$-$C_{15}$-stretching vibration was found to be located at simirlarily low wavenumber (1154 $cm^{-1}$) which indicated a 13-*cis*, 14-s-*cis* isomer (12). However, since the $C_{14}$-$C_{15}$ HC-CH torsion angle was determined to be 152 ° in $P_{480}$, this conclusion is not valid for ChR2 (3)

**Supplementary Note 5: Retinal C=N-conformation of $P_{480}$ and $D_{470}$.** The $D_2O$-induced blue shifts of the $C_{14}$-$C_{15}$-stretching vibration are indicative for the isomeric state of the C=N-Bond (13, 14). Large upshifts (>20 $cm^{-1}$) indicate a C=N-*syn* conformation whereas relatively small upshifts (5-10 $cm^{-1}$) indicate a C=N-*anti* conformation. In Fig. 3B the $P_{480}$-minus-$D_{470}$ FTIR difference spectra are shown for undeuterated (black) and deuterated samples (green). As the $C_{14}$-$C_{15}$-stretching vibration of $D_{470}$ at 1186 $cm^{-1}$ is upshifted by only +5 $cm^{-1}$ in $D_20$, the C=N-bond of the RSB in $D_{470}$ is most likely in the C=N-*anti* conformation. This is in line with NMR data of the non-photolysed state (2, 3). In $P_{480}$, the blue shift of the $C_{14}$-$C_{15}$ band at 1154 $cm^{-1}$ is around +26 $cm^{-1}$ which indicates a C=N-*syn* conformation of the RSB. In addition, the $C_{10}$-$C_{11}$ band at 1176 $cm^{-1}$ is upshifted to 1210 $cm^{-1}$, which is most likely due to delocalisation of the $C_{10}$-$C_{11}$ stretching mode (A normal mode that contains high content of other C-C-vibrations) (13). The strong coupling of the $C_{14}$-$C_{15}$ stretch to the N-H rocking vibration in $P_{480}$ further supports, that $P_{480}$ is the 13-*cis*,C=N-*syn* component of the apparent $D_{app}$ state found in a resonance Raman study (2). Because the $C_{14}$-$C_{15}$-stretching vibration band of $P_{480}$ is located at quite low wavenumbers, it is unlikely to overlap strongly with other retinal bands and therefore it can be used as a marker band for the 13-*cis*,C=N-*syn* conformation of $P_{480}$ in time resolved FTIR measurements.

**Supplementary Note 6: Kinetic behavior at different flash frequencies.** The decay of $P_{520}^{N}$ occurs with a half time of approximately 30 ms if only $D_{470}$ is excited(6). Upon increased flash frequency or continuous illumination with increasing power $P_{480}$ accumulates due to the slow decay of approx. 40 s (4). In the Raman experiments (Fig. S2B and S3-4) a maximum accumulation of 50 %-60 % was observed which can only be explained by a light induced photo reaction of $P_{480}$ which leads to a faster buildup of the $D_{470}$ (shortcut reaction). To monitor this back reaction, we compared global fit results of fastscan measurements on ChR2 with low (0.005 Hz) and high flash repetition rates (0.2 Hz). In Fig. S8 decay associated amplitude spectra of the open state decay are shown. If the repetition rate is low (0.005 Hz, black spectra), mainly $D_{470}$ is excited by the flash and the decay is predominantly described by the component with $t_{1/2}$=30 ms. Only a small fraction of a $t_{1/2}$=250 ms component is needed to describe the data properly. If the flash frequency is increased to 0.2 Hz (red spectra) a similar $t_{1/2}$=30 ms component is found, which is 50 % to 60 % less intense (the spectra are scaled by a factor of 2 for direct comparison). This can be rationalized by the fact that the amount of $D_{470}$ is decreased of about the same value in the sample. The second component with a half time of about 250 ms is drastically increased, when high frequency conditions are applied and $P_{480}$ is accumulated in the ground state. Therefore we consider the 250 ms component as the decay of the photo product of $P_{480}$. This component is also visible in the continuous light experiments (see Fig. S2).
All the decay associated amplitude spectra exhibit the negative $D_{470}$ bands at 1244 $cm^{-1}$ and 1234 $cm^{-1}$. Therefore ground state repopulation occurs with both decays. This further supports that $P_{520}^{N}$ is directly converted back to the ground state and not to $P_{480}$. By light-induced reaction of $P_{480}$ the ground state transition of $P_{480}$ is accelerated. Since the protonation state of E90 differs in $P_{480}$ and $D_{470}$ an E90 band should be visible in the 250 ms component. In fact a negative E90 band is visible in the carbonyl region of the slow component which is best visible in the H134R variant (insets, blue spectra) which has a more pronounced E90 band than the WT. Further the C=N-*syn*-marker band at 1154 $cm^{-1}$ is visible in the slow component suggesting a parallel back reaction to $P_{480}$. The slow 250 ms decay component is also visible after switching off continuous light (see Fig.S2C and S2D) in FTIR and Raman experiments (see also Fig. S9).

The two decay components are also visible in the measurements of the $^{13}C_{14}$-$^{13}C_{15}$-labelled sample. The isotopic shifts in both components exhibit similarities but are not identical. The presence of the negative $D_{470}$ band at 1234 $cm^{-1}$ in the decay associated spectra of both the $O_1$ and $O_2$ decay indicates direct buildup of $D_{470}$. The downshift of the negative $C_{14}$-$C_{15}$ band of $D_{470}$ around 1185 $cm^{-1}$ to 1170 $cm^{-1}$ dominates the double difference spectra. Therefore it is difficult to properly assign the $C_{14}$-$C_{15}$ stretch vibrations of the putative open states. However, for $O_1$ the shoulder at 1161 $cm^{-1}$ seems to be downshifted to 1145 $cm^{-1}$ and is therefore tentatively assigned to the $C_{14}$-$C_{15}$ stretch vibration of $O_1$. The effect on the positive 1170 $cm^{-1}$ band is most likely only due to the overlap with the intense $D_{470}$ band in the labeled sample. For $O_2$ no conclusion regarding C-C-stretch assignments can be drawn. The downshift of the negative $P_{480}$ band at 1154 $cm^{-1}$ that is only visible in the $O_2$ decay associated spectrum indicates that $O_2$ decays partially to $P_{480}$.

**Supplementary Note 7: Theoretical characterization of the anti- and syn-photocycle intermediates.** Upon all-*trans*,C=N-*anti* → 13-*cis*,C=N-*syn* double isomerization

($D_{470}$→$P_{480}$), the position of the RSB proton is only slightly changed, resulting in a structurally similar state and only a slightly red-shifted absorption maximum (2, 6). In contrast, single isomerization around the $C_{13}$=$C_{14}$-bond ($D_{470}$ → $P_{390a}^{M1}$) induces an upward orientation of the RSB proton (modeled structure in Fig. 5A).
For all-*trans*,C=N-*anti*→13-*cis*,C=N-*anti* isomerization we observe the same new orientation of protonated E90 as in Hummer et al. (15), after protonation of the counter ion D253 (Fig. S12-S13C+D). This new arrangement allows only single water molecules influx in the protonated state of E90, because the retinal is under tension in the anti-conformation (15). But it takes microsecond simulation times to see this water influx.
We also simulated the 13-*cis,*C=N-*syn* isomerization of $P_{480}$ with and without E90 deprotonation. Although protonation of E90 is artificial for $P_{480}$ in the WT protein, it helps to understand the role E90 deprotonation during light adaptation. In a first step, we performed the double isomerization by stepwise rotating around the $C_{15}$=N and $C_{13}$=$C_{14}$-double bonds. As long as E90 remains protonated Helix 2, 7 stays connected via E90 and D253 (Fig. 5B ; Fig. S13B). Deprotonation of E90 leads to an alternative contact between E90 and K93 (Fig. S12). This opens the central gate, results in an influx of water molecules into the pore (Fig. 5B) and causes an outward movement of Helix 2 which is in line with ESR measurements on WT-like variants (16).
Most importantly, in all the simulated structures, the inner gate remains closed (Fig. S14) and neither sodium nor proton conductance are expected to occur in the structures of $P_{480}$ and $P_{390a}^{M1}$ which represent either a closed or a pre-open photocycle intermediate.

### FTIR and Raman Measurements

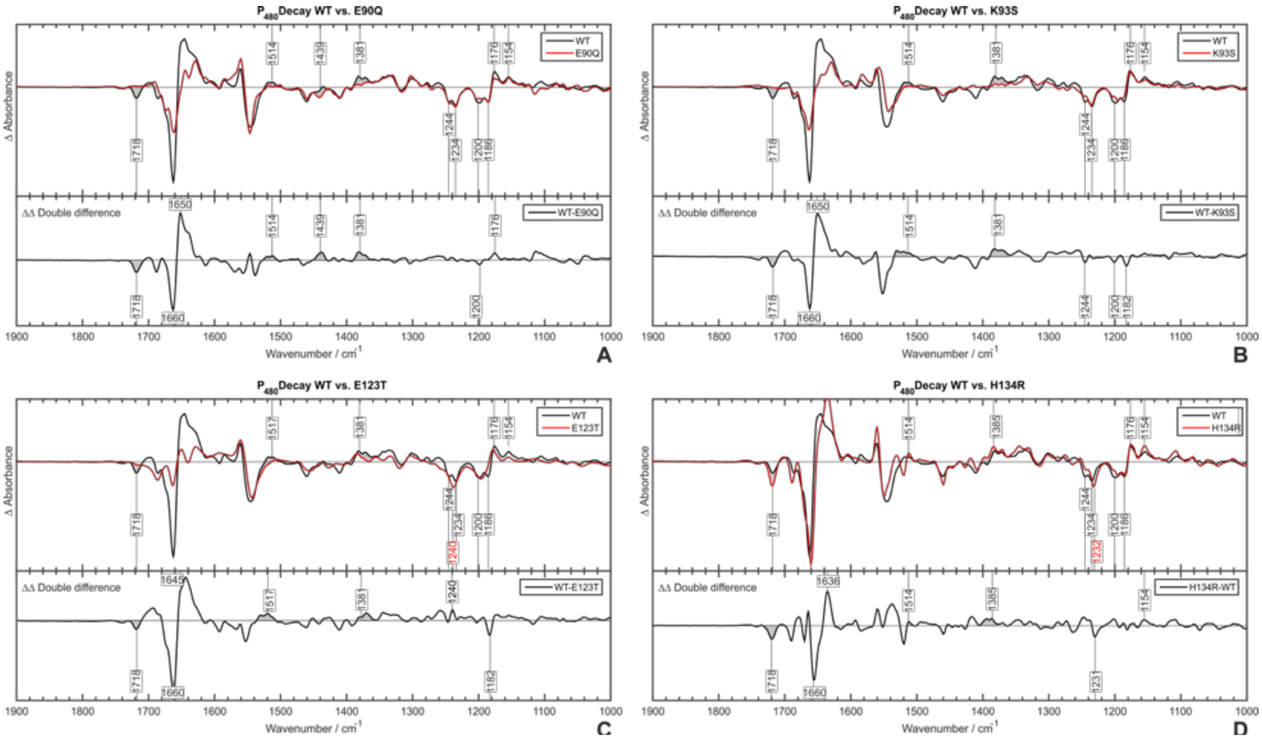


**Figure S1:** Comparison $P_{480}$-minus-$D_{470}$ FTIR difference spectra for ChR2 WT and ChR2 variants that prevent E90 deprotonation. The helix hydration marker band is reduced if no E90 deprotonation is observed.

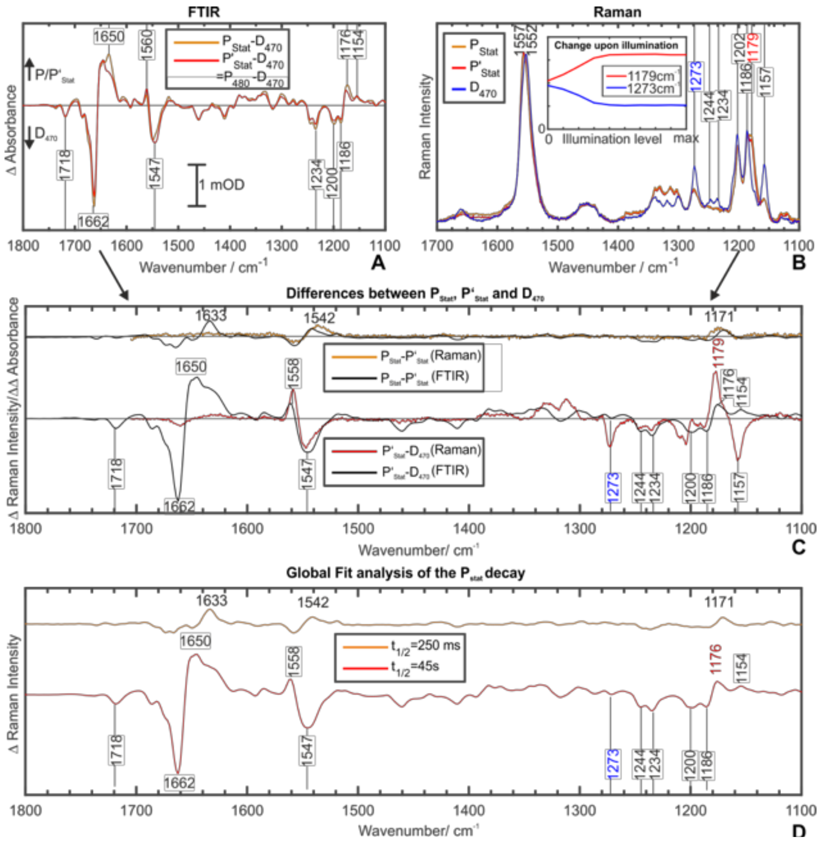


**Figure S2:** Dark state ($D_{470}$) and photostationary state measured under continuous illumination ($P_{Stat}$) and 2 s after switching off the light ($P'_{Stat}$) monitored by FTIR difference spectroscopy and pre-resonance Raman spectroscopy. (A) Only minor differences can be seen in the $P_{Stat}$-minus-$D_{470}$ (orange) and $P'_{Stat}$-minus-$D_{470}$ (red) difference spectra. The coorsponds exactly to the $P_{480}$-$D_{470}$ difference spectrum. (B) Raman spectra of $P_{Stat}$ (orange), $P'_{Stat}$ (red) and $D_{470}$ (black). Merely small differences are visible in the spectra of $P_{Stat}$ (orange) and $P'_{Stat}$ (red) as well. However both spectra differ strongly from the spectrum of $D_{470}$ (blue). The inset shows the change of the $P_{Stat}$ and $D_{470}$ marker bands (1179 cm$^{-1}$ and 1273 cm$^{-1}$) depending on the illumination intensity. (C) The Raman $P_{Stat}$-minus-$P'_{Stat}$ (orange) and $P'_{Stat}$-minus-$D_{470}$ (red) difference spectra correspond to the respective difference spectra measured by FTIR, which demonstrates that the same processes are monitored by the two spectroscopic techniques. (D) Global Fit analysis of the $P_{Stat}$ decay revealed two components. The decay associated amplitude spectra represent the fast (orange) and slow (red) decay component.

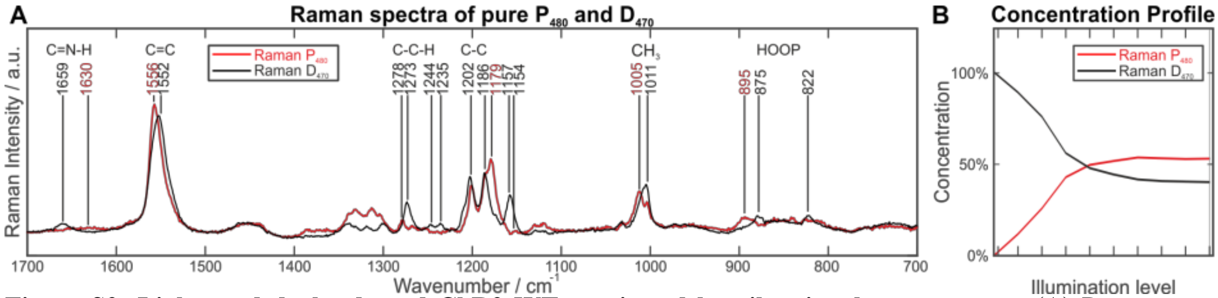


**Figure S3: Light- and dark-adapted ChR2 WT monitored by vibrational spectroscopy.** (A) Raman spectra of pure $P_{480}$ (red) and pure $D_{470}$ (black) are shown. Under photo-stationary conditions (continuous blue light) an illumination intensity-dependent mixture of both spectra is formed (B).

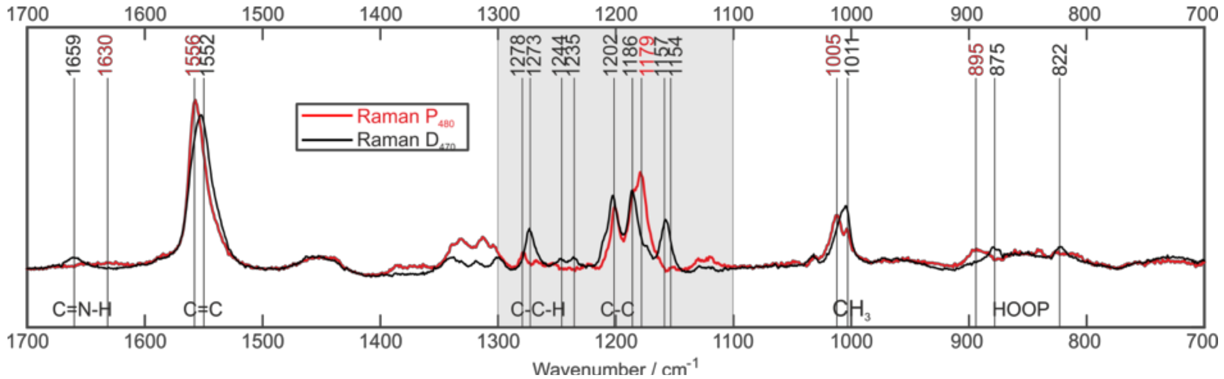

**Figure S4:** Pure P480 and D470 spectra generated by a subtraction procedure.

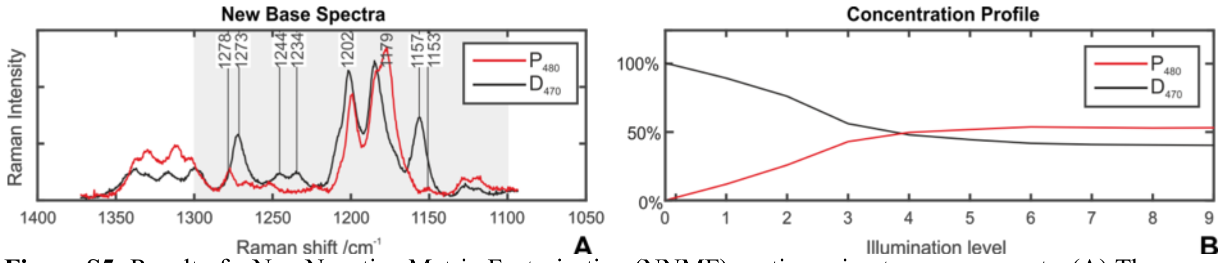


**Figure S5:** Result of a Non Negative Matrix Factorisation (NNMF) routine using two components. (A) The new base spectra represent the absolute components $D_{470}$ (black) and $P_{480}$ (red). (B) The illumination dependent concentration profiles show, that upon illumination $D_{470}$ is converted into $P_{480}$. This process saturates at higher illumination power which results in a max. turnover of approx. 50%-60%.

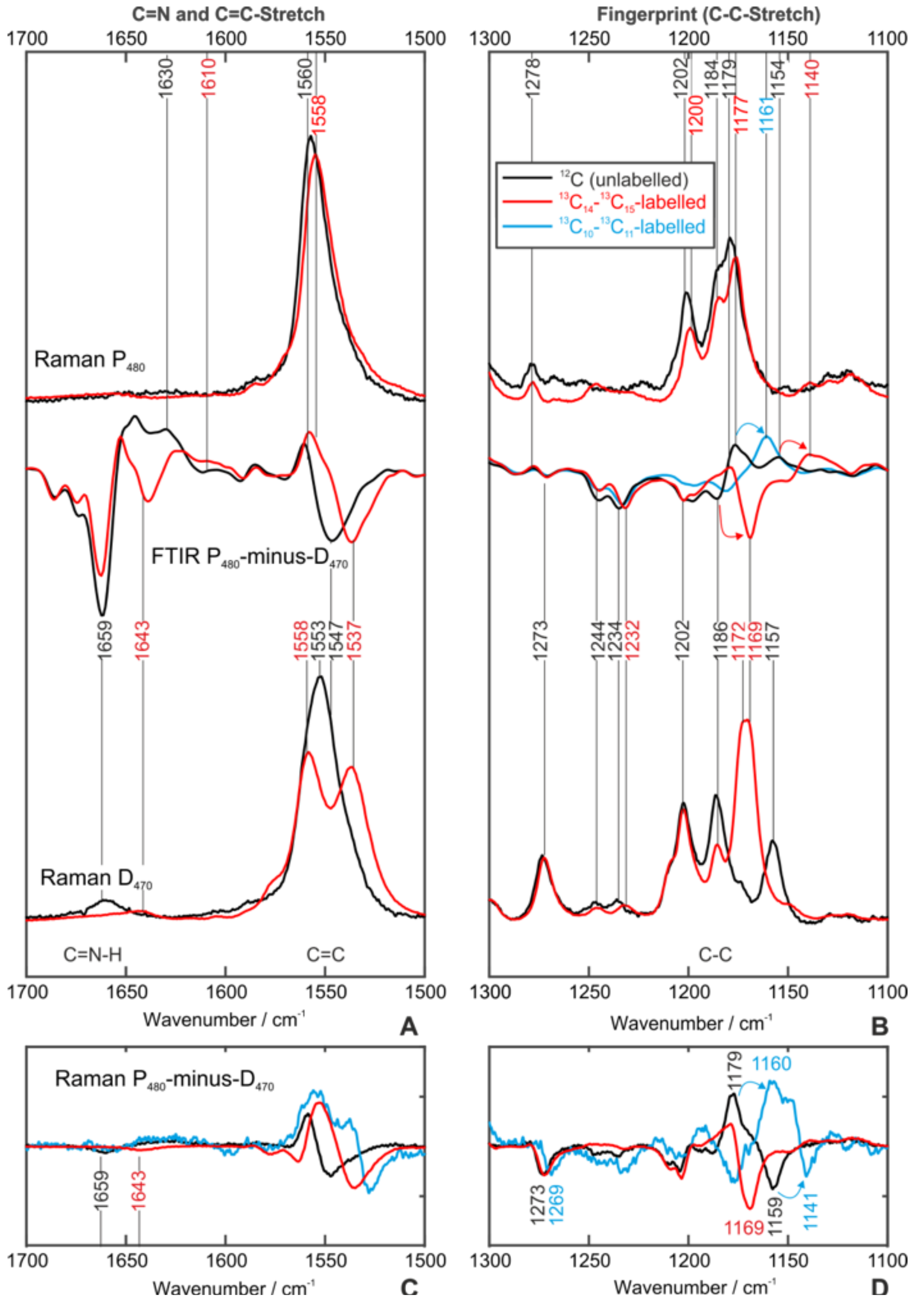


**Figure S6:** Comparison of $P_{480}$ and $D_{470}$ pre-resonance Raman spectra and $P_{480}$-minus-$D_{470}$ FTIR difference spectra for unlabeled (black) and $^{13}C_{14}$-$^{13}C_{15}$-labeled (red) samples. In the bottom Raman $P_{480}$-minus-$D_{470}$ difference spectra are compared for unlabeled (black) $^{13}C_{14}$-$^{13}C_{15}$-labelled (red) unlabeled (black) and $^{13}C_{10}$-$^{13}C_{11}$-labeled (red) samples.

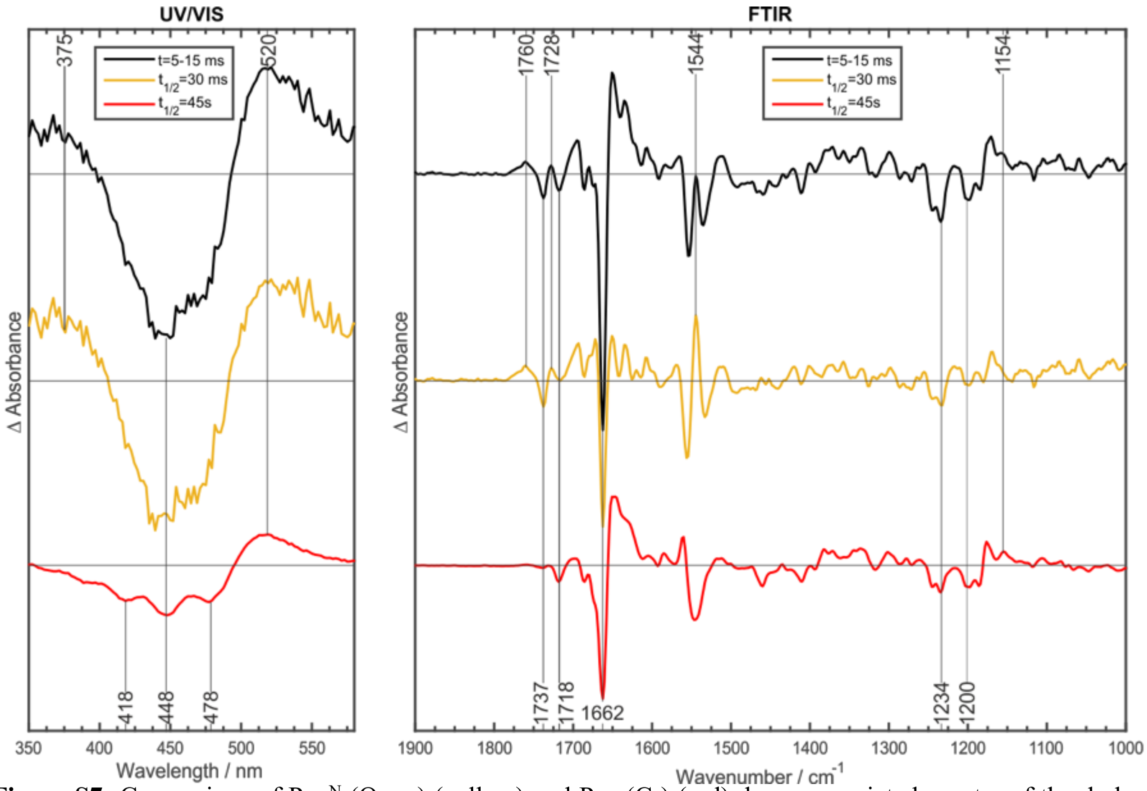


**Figure S7:** Comparison of $P_{520}^{N}$ ($O_{1\text{-late}}$) (yellow) and $P_{480}$ ($C_2$) (red) decay-associated spectra of the dark-adapted photocycle. The negative $D_{470}$ ($C_1$) bands at 1234 cm$^{-1}$ indicate the direct conversion of these species into $D_{470}$. Since their amplitude is similar, a similar efficiency of the formation of $P_{520}^{N}$ ($O_{1\text{-late}}$) and $P_{480}$ ($C_2$) can be assumed for the dark-adapted cycle. The corresponding difference spectra at 5-15 ms are shown in black.

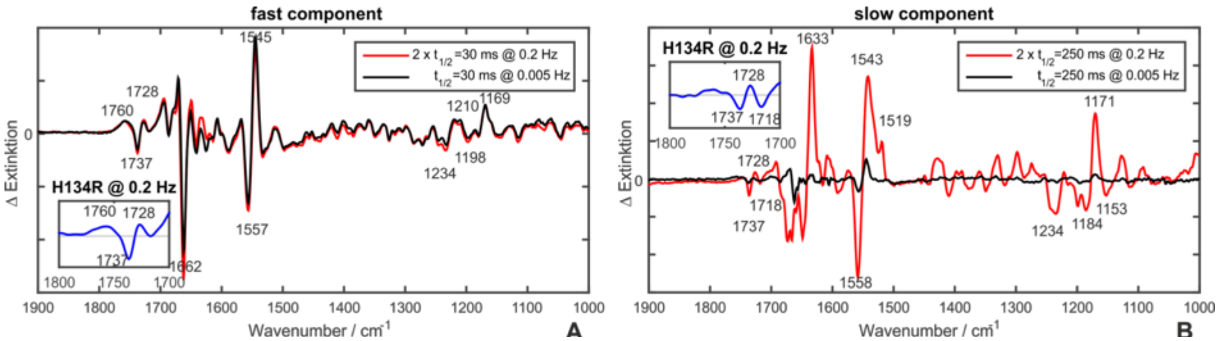


**Figure S8:** Measurements on WT and H134R with different flash repetition rates. Two components are sufficient to describe the decay of the open state. (A) Comparison of the $t_{1/2}$=30 ms component for low (0.005 Hz, black) and high (0.2 Hz,red) flash repetition rates. (B) Comparison of the $t_{1/2}$=250 ms component for low (0.005 Hz, black) and high (0.2 Hz,red) flash repetition rates. This slow decaying component gains intensity as $P_{480}$ is accumulated. This slower component matches the decay component found in the FTIR and Raman continuous illumination experiments shown in Fig. S2 and S7.

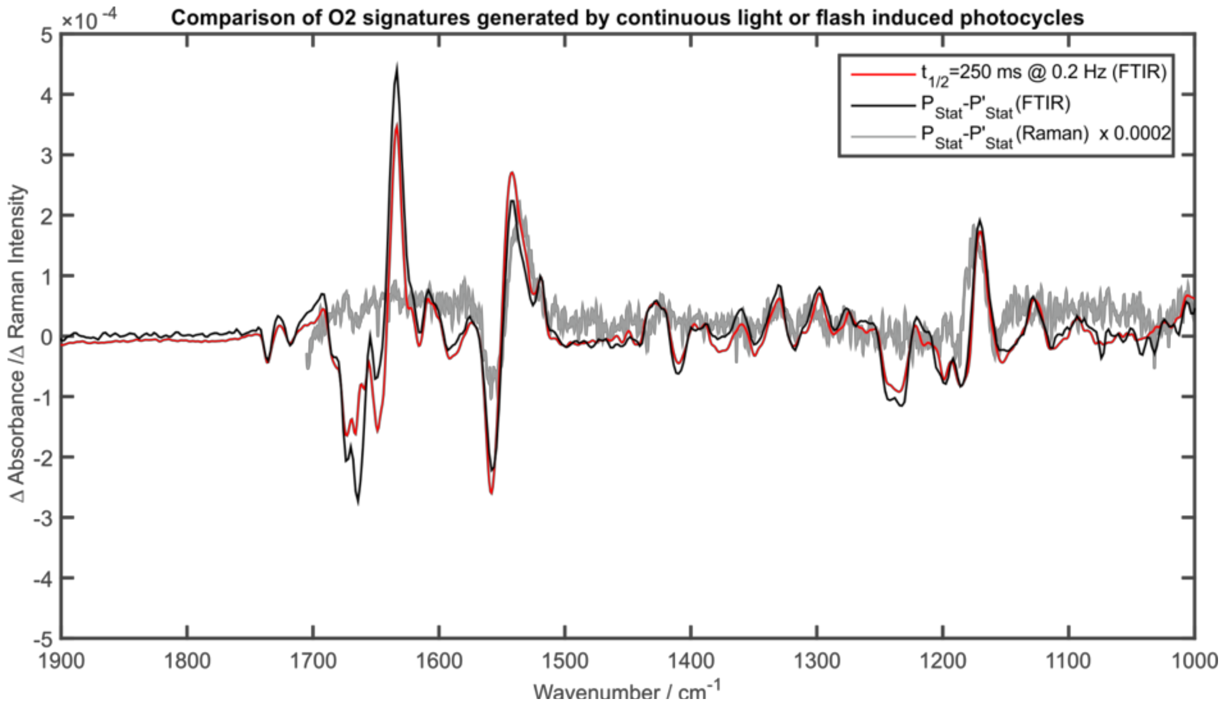


**Figure S9:** Comparison of the $O_2$ decay associated spectrum measured by FTIR with high flash repetition rates with the $P_{Stat}$-minus $P'_{Stat}$ spectra from the continuous light experiments displayed in Fig. S2.

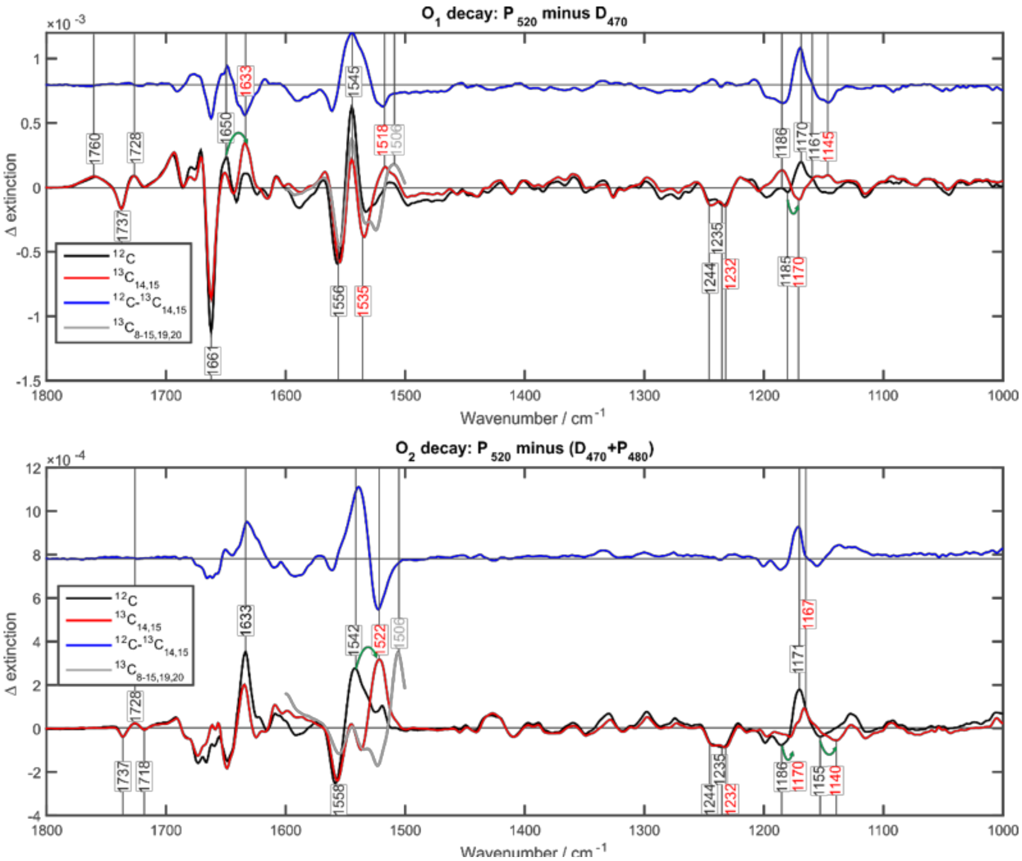


**Figure S10:** Comparison of $O_1$ and $O_2$ decay associated amplitude spectra for unlabeled (black) $^{13}C_{14}$-$^{13}C_{15}$-labelled (red) samples. In blue unlabeled-minus-labelled double difference spectra are shown.

### Electrophysiological patch clamp recordings

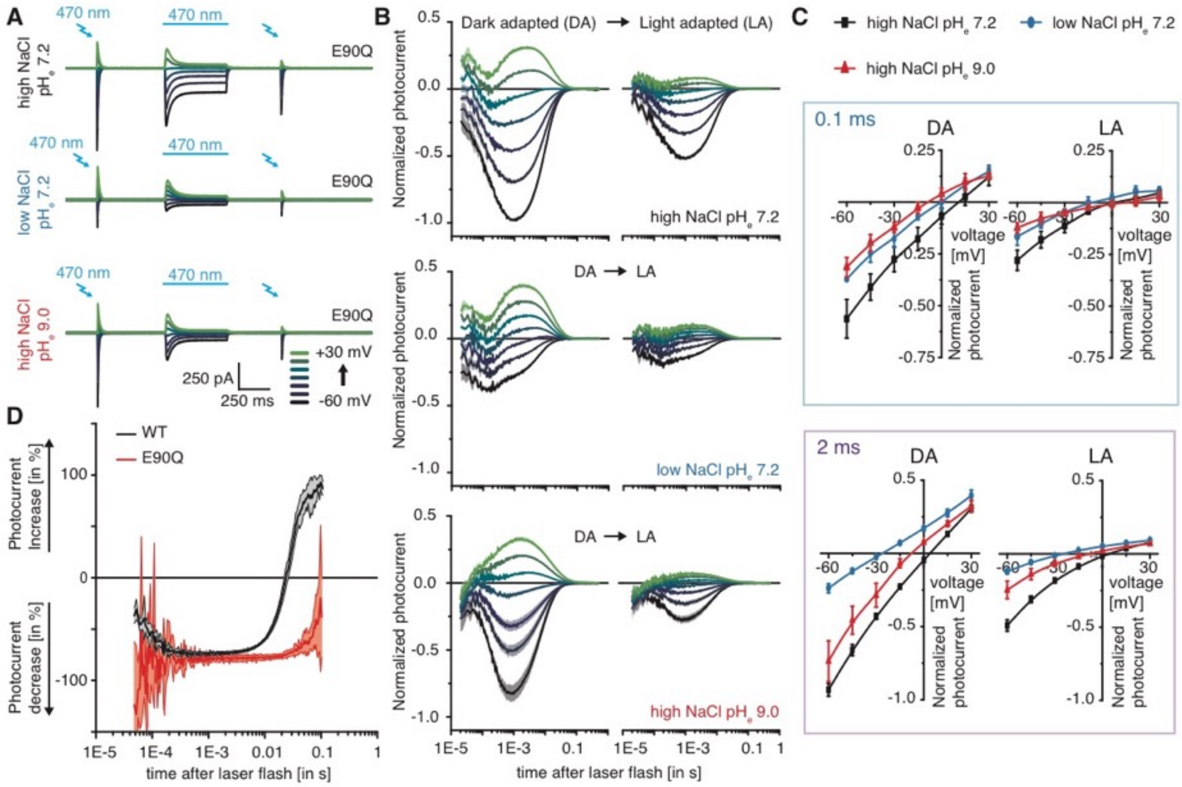


**Figure S11: Patch-clamp recording in HEK293 cells of proton and sodium conductance in dark and light adapted ChR2 E90Q.** (A) Representative photocurrents of ChR2 E90Q with intracellular 110 mM NaCl $pH_i$ 7.2 and extracellular 110 mM $Na^+$ $pH_e$ 7.2 (top), 1 mM $Na^+$ $pH_e$ 7.2 (middle), 110 mM $Na^+$ $pH_e$ 9.0 (bottom) at different holding voltages. Photocurrents were excited before and after light adaptation by 7 ns laser pulses of 470 nm light. For light adaptation cells were illuminated for 500 ms with continuous 470 nm light. (B) Normalized, log-binned and averaged photocurrents of the dark adapted (DA) or light adapted (LA) E90Q mutant in the different buffering conditions shown in A (mean ± SE, n=5-6). (C) Current-voltage dependency of normalized photocurrents of ChR2 E90Q 0.1 ms (top) and 2 ms (bottom) after excitation in different extracellular buffer compositions before (DA) and after (LA) light adaptation (Mean ± SD, n=5-6) (D) Relative photocurrent change upon light adaptation of ChR2 WT and ChR2 E90Q at different extracellular voltages and $pH_e$ (I(LA)-I(DA))/I(DA), mean ± SE, n=5)

### MD Simulations

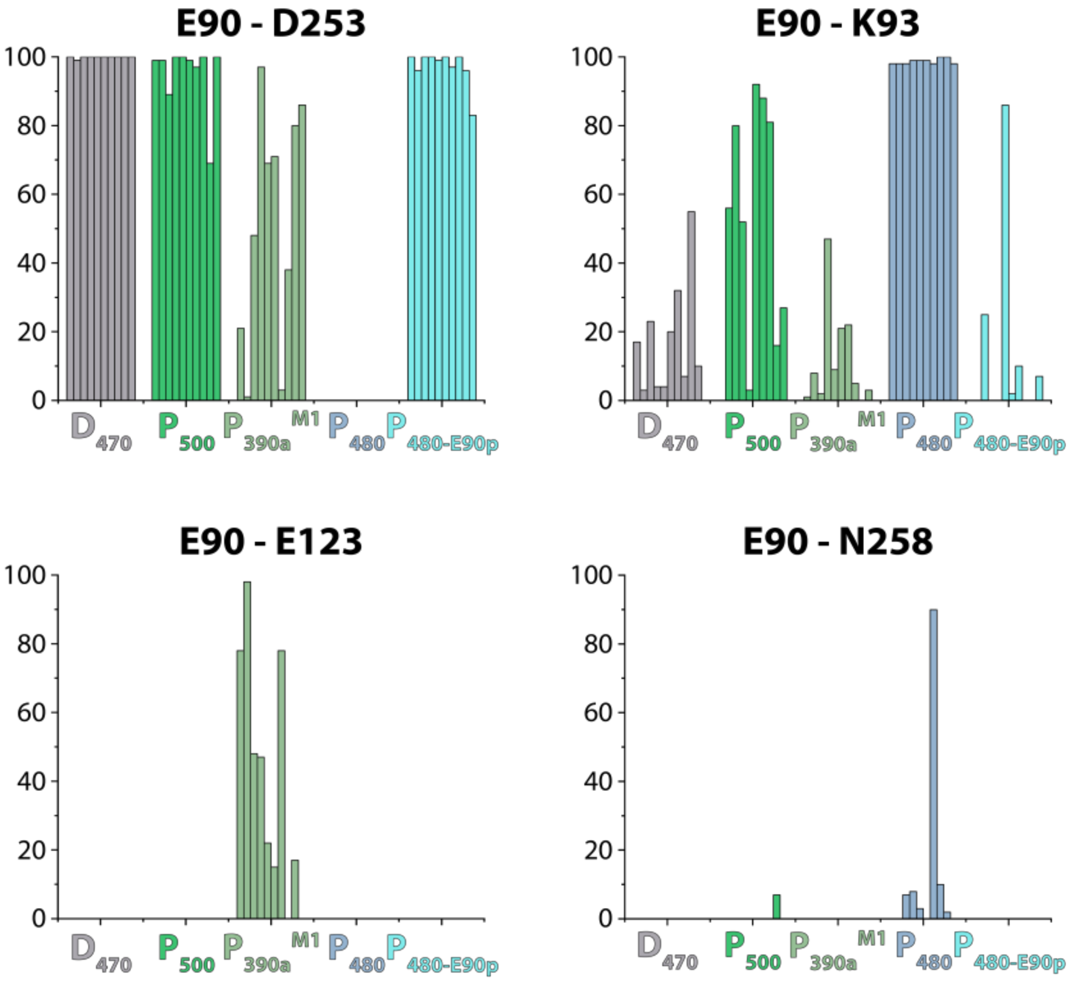


**Figure S12:** Overview of E90 hydrogen bond pattern for 5 independent simulations with 2 monomers forming 1 dimer based on the Channelrhodopsin 2 crystal structure (PDB-ID 6eid (17)). $D_{470}$ (all-*trans*,C=N-*anti*, E90p) represents the ground state $C_1$. $P_{500}$ illustrates the single isomerized state of the anti-photocycle. In a next step the RSB was deprotonated and D253 was protonated ($P_{390a}^{M1}$). In a second approach the syn-photocycle was calculated. $P_{480}$ shows the double isomerized state of the light adapted ground state $C_2$ (13-*cis*,C=N-*syn*, E90dp). $P_{480}$-E90P shows an artificial state of $P_{480}$ with protonated E90 after double isomerization.

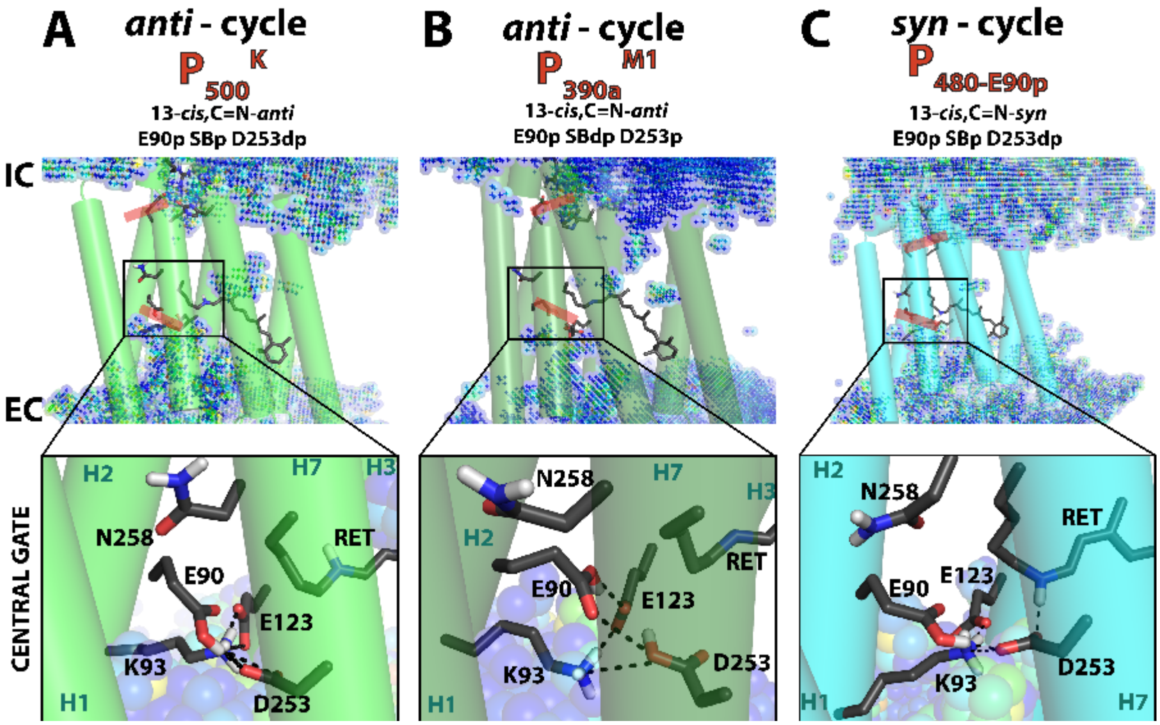


**Figure S13: Formation of *anti*- and *syn*-photocycle intermediates.** The representative structures of the intermediates based on Channelrhodopsin 2 crystal structure (PDB-ID 6eid(17)) are shown. (A) The panel shows an artificial state in the *syn*-photocycle after the all-*trans*,C=N-*anti* → 13-*cis*,C=N-*syn* double isomerization without E90 deprotonation ($P_{480}$-E90p). (B) The panel shows the structure of the intermediate $P_{500}{}^{K}$ in the *anti*-photocycle containing 13-*cis*,C=N-*anti* retinal after single isomerization. (C) The panel shows the structure of the intermediate $P_{390a}{}^{M1}$ in the dark-adapted photocycle containing 13-*cis*,C=N-*anti* retinal after single isomerization, RSB deprotonation and D253 protonation.

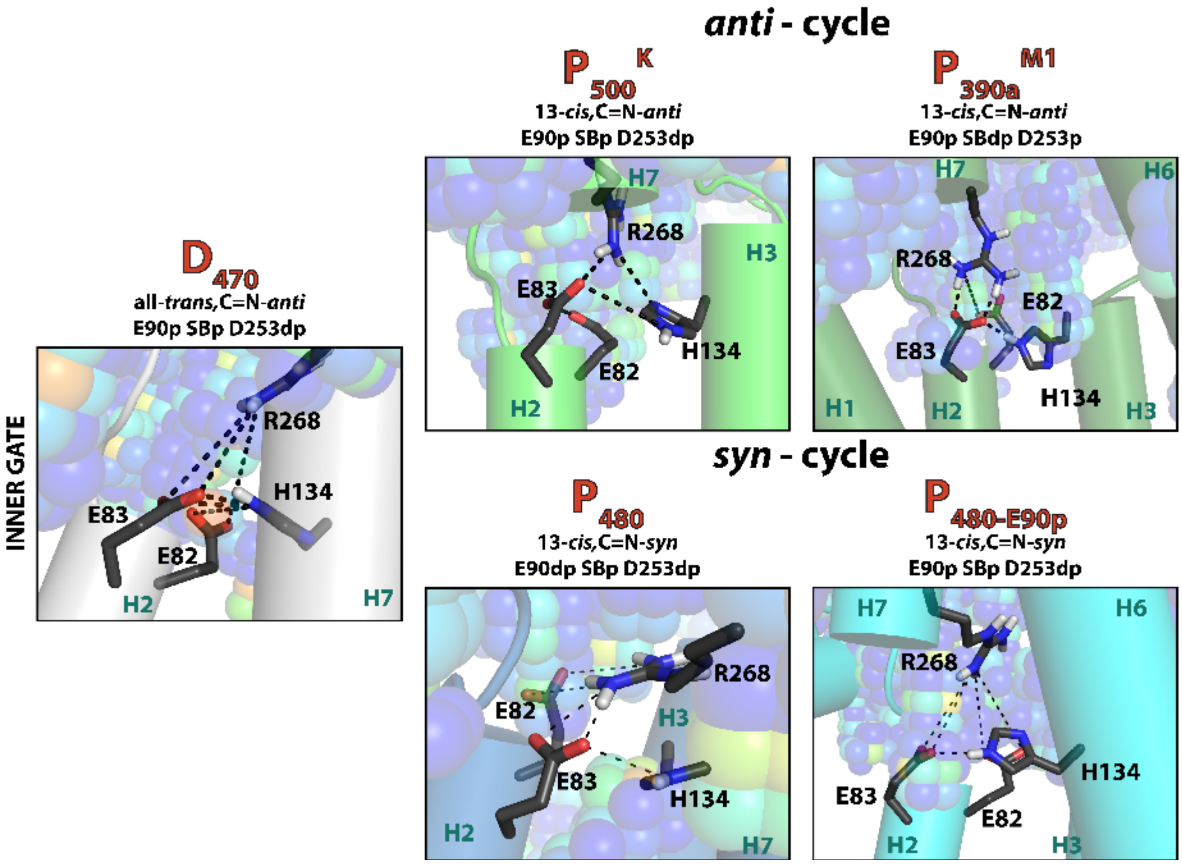


**Figure S14: Inner gate interactions in the *anti*- and *syn*-phtocycle.** The representative structures for the inner gate of all calculated intermediates on Channelrhodopsin 2 crystal structure (PDB-ID 6eid (17)) are shown. In all simulated structures the inner gate remains closed.

**Table S1: Assignment of C-C, and C=N stretching modes.** The numbers in parenthesis indicate wavenumber shift upon retinal $^{13}C_{14}$-$^{13}C_{15}$ or $^{13}C_{10}$-$^{13}C_{11}$ isotopic labelling respectively. For a detailed discussion see Fig. S3.

| vibrational modes | **$D_{470}$** | | **$P_{480}$** | |
|---|---|---|---|---|
| | Raman | IR | Raman | IR |
| **$C_{14}$-$C_{15}$**<br>$D_2O$-shift | 1186 (-17)<br>+8 | 1186 (-17)<br>+5 | 1154 (-14)<br>- | 1154 (-14)<br>+26 |
| **$C_{10}$-$C_{11}$**<br>$D_2O$-shift | 1157 (-16)<br>0 | - | 1179 (-18)<br>+22 | 1176 (-15)<br>+22 |
| **$^{12}C$=N**<br>**$^{13}C$=N** | 1659<br>1643 | -<br>1643 | 1630<br>- | 1630<br>1610 |

### References


1. The retinal structure of channelrhodopsin-2 assessed by resonance Raman spectroscopy. - PubMed - NCBI Available at: http://www.ncbi.nlm.nih.gov/pubmed/19854176 [Accessed March 2, 2016].

2. Bruun S, et al. (2015) Light–Dark Adaptation of Channelrhodopsin Involves Photoconversion between the all- *trans* and 13- *cis* Retinal Isomers. *Biochemistry* 54(35):5389–5400.

3. Becker-Baldus J, et al. (2015) Enlightening the photoactive site of channelrhodopsin-2 by DNP-enhanced solid-state NMR spectroscopy. *Proceedings of the National Academy of Sciences* 112(32):9896–9901.

4. Ritter E, Stehfest K, Berndt A, Hegemann P, Bartl FJ (2008) Monitoring Light-induced Structural Changes of Channelrhodopsin-2 by UV-visible and Fourier Transform Infrared Spectroscopy. *J Biol Chem* 283(50):35033–35041.

5. Kuhne J, et al. (2015) Early Formation of the Ion-Conducting Pore in Channelrhodopsin-2. *Angew Chem Int Ed* 54(16):4953–4957.

6. Eisenhauer K, et al. (2012) In channelrhodopsin-2 Glu-90 is crucial for ion selectivity and is deprotonated during the photocycle. *J Biol Chem* 287(9):6904–6911.

7. Lórenz-Fonfría VA, et al. (2015) Temporal evolution of helix hydration in a light-gated ion channel correlates with ion conductance. *Proc Natl Acad Sci USA* 112(43):E5796-5804.

8. Aton B, Doukas AG, Callender RH, Becher B, Ebrey TG (1977) Resonance Raman studies of the purple membrane. *Biochemistry* 16(13):2995–2999.

9. Berry MW, Browne M, Langville AN, Pauca VP, Plemmons RJ (2007) Algorithms and applications for approximate nonnegative matrix factorization. *Computational Statistics & Data Analysis* 52(1):155–173.

10. Smith SO, et al. (1985) Vibrational analysis of the all-trans retinal protonated Schiff base. *Biophys J* 47(5):653–664.

11. Smith SO, et al. (1987) Vibrational analysis of the all-trans-retinal chromophore in light-adapted bacteriorhodopsin. *Journal of the American Chemical Society* 109(10):3108–3125.

12. Gerwert K, Siebert F (1986) Evidence for light-induced 13-cis, 14-s-cis isomerization in bacteriorhodopsin obtained by FTIR difference spectroscopy using isotopically labelled retinals. *EMBO J* 5(4):805–811.

13. Smith SO, et al. (1984) Determination of retinal Schiff base configuration in bacteriorhodopsin. *PNAS* 81(7):2055–2059.

14. Vogel R, et al. (2003) Deactivation of Rhodopsin in the Transition from the Signaling State Meta II to Meta III Involves a Thermal Isomerization of the Retinal Chromophore CN Double Bond. *Biochemistry* 42(33):9863–9874.

15. Ardevol A, Hummer G (2018) Retinal isomerization and water-pore formation in channelrhodopsin-2. *PNAS* 115(14):3557–3562.

16. Sattig T, Rickert C, Bamberg E, Steinhoff H-J, Bamann C (2013) Light-induced movement of the transmembrane helix B in channelrhodopsin-2. *Angew Chem Int Ed Engl* 52(37):9705–9708.

17. Volkov O, et al. (2017) Structural insights into ion conduction by channelrhodopsin 2. *Science* 358(6366):eaan8862.
